# A microbiota-derived bile acid overcomes antibiotic-induced hyporesponsiveness to immune checkpoint therapy by enhancing CD8^+^ T cell antitumor immunity

**DOI:** 10.64898/2026.04.15.718788

**Authors:** Wenling Li, Christina M. Zarek, Hesuiyuan Wang, Shuheng Gan, Parastoo Sabaiefard, Priscilla Del Valle, Jiwoong Kim, Nicole Poulides, Laura A. Coughlin, Jake N. Lichterman, Cheng Zhang, Rebecca Suet-Yan Chiu, Tarun Srinivasan, Mauricio J. Velasquez, Indu Raman, Vanessa J. Maddox, Jeffrey McDonald, Ralf Kittler, Prithvi Raj, Xin V. Li, Xiaowei Zhan, Chen Liao, Joao B. Xavier, Andrew Y. Koh

**Affiliations:** Department of Pediatrics, The University of Texas Southwestern Medical Center, Dallas, TX, USA; Department of Population and Data Sciences, The University of Texas Southwestern Medical Center, Dallas, TX, USA; Department of Internal Medicine, The University of Texas Southwestern Medical Center; Dallas, TX, USA; Department of Pathology, The University of Texas Southwestern Medical Center; Dallas, TX, USA; Department of Immunology, The University of Texas Southwestern Medical Center; Dallas, TX, USA; Center for Human Nutrition, The University of Texas Southwestern Medical Center; Dallas, TX, USA; McDermott Center for Human Growth and Development, The University of Texas Southwestern Medical Center; Dallas, TX, USA; Program for Computational and Systems Biology, Memorial Sloan Kettering Cancer Center, New York, NY, USA; Department of Microbiology and Immunology, Geisel School of Medicine, Dartmouth, Hanover, NH, USA; Department of Microbiology, The University of Texas Southwestern Medical Center, Dallas, TX 75390; Harold C. Simmons Comprehensive Cancer Center, The University of Texas Southwestern Medical Center, Dallas, TX 75390

## Abstract

Gut microbiota are critical determinants of effective immune checkpoint therapy (ICT), yet the microbial mediators and host mechanisms that enhance antitumor immunity remain poorly understood. Here, we identify the microbiota-derived bile acid taurodeoxycholic acid (TDCA) as a metabolite associated with immune checkpoint therapy (ICT) response. TDCA administration alone is sufficient to overcome antibiotic-induced ICT hyporesponsiveness across multiple murine tumor models. Mechanistically, TDCA directly enhances CD8⁺ T cell–mediated antitumor immunity, increasing cytotoxicity. These effects required signaling through the bile acid receptor TGR5. Together, these findings reveal TDCA as a gut microbial metabolite that restores ICT efficacy after antibiotic disruption by directly augmenting CD8⁺ T cell anti-tumor activity. This work supports metabolite replacement as a therapeutic strategy to mitigate antibiotic-associated loss of cancer immunotherapy response.

**Significance:** TDCA is a microbiota-derived metabolite that restores immune checkpoint therapy efficacy after antibiotic disruption by directly enhancing CD8⁺ T-cell–mediated anti-tumor immunity through bile acid receptor TGR5 signaling. Our findings suggest that supplementation with defined microbial metabolites can mitigate antibiotic-associated loss of immunotherapy response without requiring broader microbiome reconstitution.

## Introduction

Immune checkpoint blockade has reshaped cancer treatment for a subset of patients by restoring dysfunctional T-cell responses and enabling durable tumor control (1,2). Most cancer patients, however, do not respond to immune checkpoint inhibitor therapy (ICT), highlighting the need to identify modifiable factors that determine therapeutic responsiveness (3,4). In recent years, the intestinal microbiota has emerged as a critical determinant of antitumor immunity and response to ICT, through both preclinical and clinical studies (5–13). Indeed, antibiotic-induced disruption of the gut microbiota has been associated with impaired responses to ICT and reduced survival in multiple cancer types (13–15). Yet antibiotics are commonly prescribed during cancer treatment because patients are often immunocompromised and vulnerable to infectious complications. Understanding the mechanisms that drive antibiotic-induced ICT hyporesponsiveness, and how this loss of response might be mitigated, remains an important unmet need.

Antibiotic treatment disrupts microbial communities and their metabolites, which shape host immunity (16). Among these, bile acids have emerged as immunomodulators capable of influencing both innate and adaptive immune responses (17–23). Bile acids are cholesterol-derived molecules that can be classified into primary and secondary bile acids (24–26). Primary bile acids are synthesized in the liver and subsequently modified by the gut microbiota into secondary bile acids (24–26), making the bile acid pool highly sensitive to microbiota changes. Accordingly, antibiotics and other microbiota-modulating agents can reshape bile acid composition and, in turn, alter downstream host immune signaling pathways (25,27).

Bile acids can either inhibit or enhance antitumor immunity – depending on the specific metabolite, receptor, and tumor setting. Deoxycholic acid (DCA) suppresses CD8⁺ T cell effector function in colorectal cancer, promoting tumor growth (21). Similarly, accumulated conjugated primary bile acids and lithocholic acid conjugates impair CD8^+^ T-cell survival and function in liver tumors (23). In contrast, ursodeoxycholic acid supplementation promoted antitumor immunity in liver cancer (23), and microbiota-conjugated secondary bile acid isomers enhanced ICT efficacy in bladder cancer by promoting a stem-like CD8^+^ T cell state (28). Collectively, these studies support a context-dependent role for bile acids in cancer immunity. But how microbiota-derived bile acids influence antibiotic-associated loss of ICT responsiveness remains unclear.

Here, we investigated how antibiotic-induced disruption of the gut microbiota alters melanoma tumor control during ICT. Using integrated metabolomic and functional analyses, we identify a microbiota-derived secondary bile acid, taurodeoxycholic acid (TDCA), as a determinant of ICT responsiveness. We demonstrate that TDCA treatment is sufficient to overcome antibiotic-induced ICT responsiveness through a CD8^+^ T cell-dependent mechanism. TDCA enhances CD8^+^ T cell effector function and cytotoxicity through the bile acid receptor TGR5. These findings identify a microbiota-derived bile acid–TRG5–CD8^+^ T cell axis as a mechanism linking antibiotic-induced microbiome disruption to impaired immunotherapy response, with potential translational relevance for cancer immunotherapy treatment.

## Results

### Antibiotic perturbation and unbiased metabolomic profiling identify TDCA as a determinant of ICT responsiveness

Antibiotics are frequently administered to cancer patients and are associated with reduced immune checkpoint therapy (ICT) efficacy in clinical studies, a finding recapitulated in preclinical models (6,14). Yet, the mechanisms linking antibiotic-induced microbiome disruption to impaired antitumor immunity remain poorly understood. Thus, we used six clinically relevant antibiotics (penicillin/streptomycin, vancomycin, levofloxacin, piperacillin/tazobactam, clindamycin, or cefepime) to generate distinct gut microbiome and metabolome states in the B16-F10 melanoma model with ICT. C57BL/6J mice were treated for one week with an antibiotic before subcutaneous implantation of B16-F10 melanoma cells. Four days after tumor inoculation, mice received α-PD-1 and α-CTLA-4 (ICT) every four days, and tumor growth was monitored longitudinally. Stool, cecal content, serum, and tumor samples were collected on day 17 post tumor inoculation (**Fig. 1A, Fig. S1A**).

**Figure 1.**
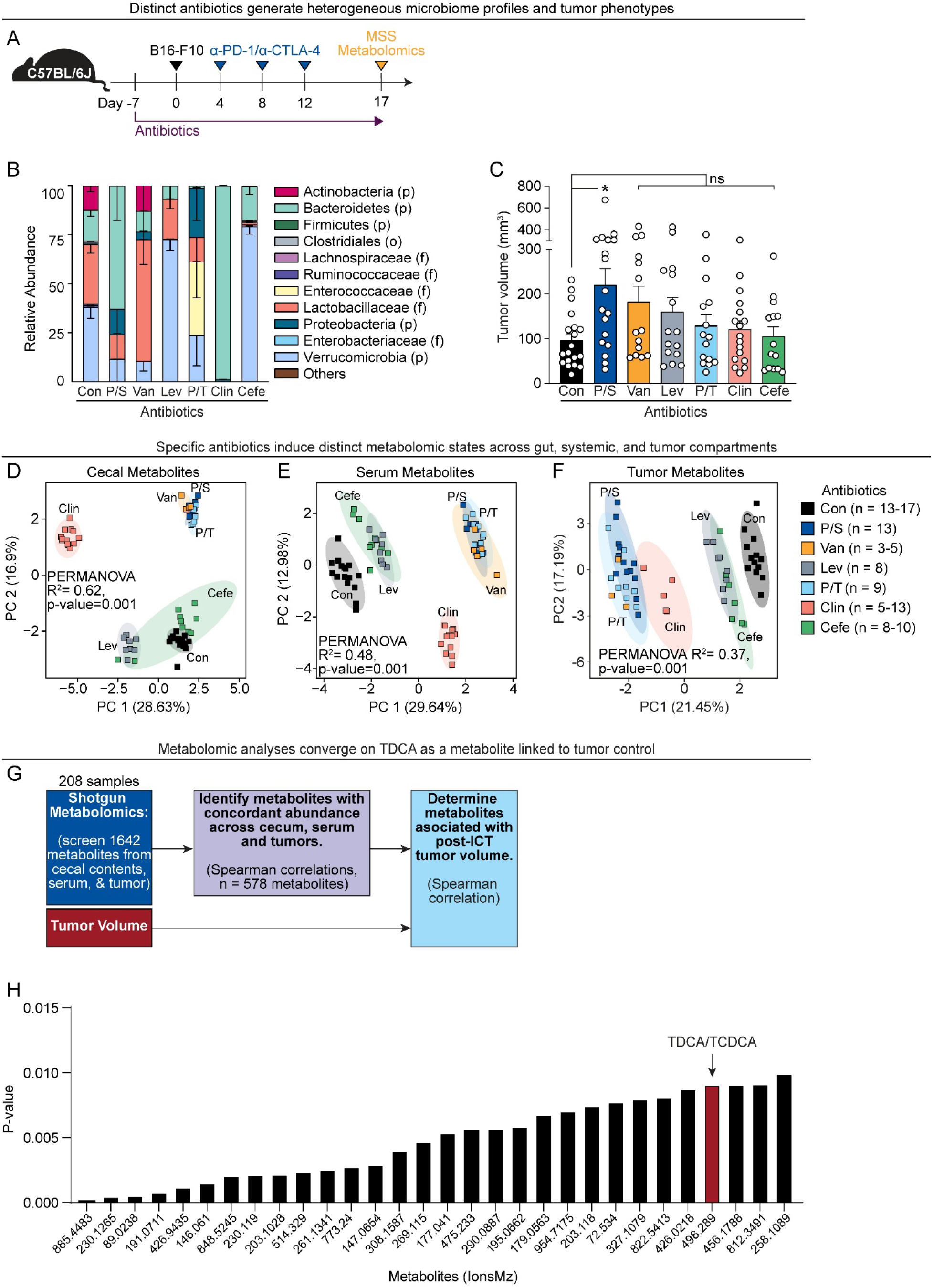
Antibiotic perturbation and unbiased metabolomic profiling identify TDCA as a determinant of ICT responsiveness. **(A)** Schematic overview of the B16-F10 heterotopic model, metabolomics and metagenomic shotgun sequencing (MSS). Mice (C57BL/6, female, 6–8 weeks old) were treated with different antibiotics throughout the experiment, starting one week before tumor injections: Con, no antibiotics control; P/S, Penicillin (0.95 g/L)/ streptomycin (2 g/L); Van, Vancomycin (0.5 g/L); Lev, Levofloxacin (0.5 g/L); P/T, Piperacillin/Tazobactam (12.5 mg/ml, I.P injection, once a day); Clin, Clindamycin (0.5 g/L); Cefe, Cefepime (12.5 mg/ml, I.P injection, twice a day). P/S, Van, Lev, Clin were administrated to mice via drinking water. 1 x 10^5^ B16-F10 cells were injected subcutaneously into the right inguinal flank. For ICT treatment, 200 µg of α-PD-1 and 200 µg α-CTLA4 (ICT) were injected intraperitoneally, as shown. On day 17 post tumor inoculation, cecal content, serum and tumors were collected for metabolomics and stool was collected for MSS. **(B)** Gut microbiota alterations from antibiotic treatments from (A), profiled by MSS of fecal gDNA. Relative abundances of bacterial phyla, averaged per antibiotic group. *n* = 5–17 mice per group. **(C)** Tumor volumes of ICT- and antibiotic-treated mice from (A), on day 17 post tumor inoculation. *n* = 14–19 mice per group. **(D)** Principle component analysis with PERMANOVA of cecal metabolites, from (A). *n* = 5–17 mice per group. **(E)** Principle component analysis with PERMANOVA of serum metabolites, from (A). *n* = 5–17 mice per group. **(F)** Principle component analysis with PERMANOVA of tumor metabolites, from (A). *n* = 3–13 mice per group. **(G)** Multi-omics and machine learning approach to identify metabolites associated with melanoma tumor response to ICT. **(H)** P-values of top 30 metabolites associated with tumor size post ICT treatment. The red bar represents the ion associated with TDCA/TCDCA. ICT, immune checkpoint inhibitor therapy. TCDCA, taurochenodeoxycholic acid. TDCA, taurodeoxycholic acid. (C) Statistical analysis by one-way ANOVA, Kruskal-Wallis test with multiple comparisons. Points represent individual mice. Bars denote mean ± SEM. All results are representative of ≥2 independent experiments. \**p*< 0.05; \*\****p*** < 0.005; ns, not significant.

Penicillin/streptomycin (Pen/Strep), vancomycin (Van), and piperacillin/tazobactam (P/T) significantly reduced total bacterial levels in the gut (**Fig. S1B**). Metagenomic shotgun sequencing (MSS) of feces revealed that each antibiotic induced a distinct pattern of microbial community restructuring (**Fig. 1B**), with Pen/Strep-, Van-, and clindamycin-treated groups showing altered beta-diversity (Bray-Curtis) compared to the control group and other antibiotic treatments (**Fig. S1C**). These antibiotic-induced microbiome states were associated with divergent responses to ICT: mice treated with Pen/Strep developed significantly larger tumors, indicated by increased tumor volumes and weights, compared to mice that received no antibiotics (**Fig. 1C, Fig. S1D**). Importantly, antibiotic treatment had no effect on tumor growth in mice receiving isotype control antibodies, indicating that antibiotics impair tumor control by reducing ICT responsiveness rather than by altering intrinsic tumor growth (**Fig. S1E–G**).

To identify microbiota-derived metabolites associated with ICT response, we performed untargeted liquid phase chromatography mass spectrometry (LC-MS) metabolomic profiling across tumor, cecal content, and serum samples (detecting 1,642 metabolites in 208 total samples, **Tables S1, S2**). Antibiotic treatment broadly reprogrammed metabolomes across all 3 compartments (principle component analysis with PERMANOVA, **Fig. 1D–F**). Notably, samples from Pen/Strep-, Van-, or P/T-treated mice clustered together across tissues (**Fig. 1D–F**). These same antibiotic regimens also produced the greatest depletion and restructuring of the gut microbiota, consistent with our prior observations in other settings (29,30), suggesting that antibiotics with the strongest effects on microbial community composition exert the largest downstream effects on the host metabolome. We next prioritized metabolites that had the same abundance changes across tissues (i.e., increased or decreased in all three tissues compared to control, determined by Spearman correlations; Benjamini-Hochberg, adjusted p-value < 0.05). 578 metabolites had positive correlations of abundances between all three tissues. To identify metabolites most strongly associated with tumor burden, independent of antibiotic treatment, we used a Spearman correlation between the tumor abundances of the 578 metabolites and the tumor volume (**Fig. 1G**). To focus on candidates with potential clinical relevance, we compared the top-ranked metabolites from this study (**Fig. 1H**) with our previously published clinical study (specifically metabolites associated with response to α-PD-1 and α-CTLA4 therapy in melanoma patients (31)). This analysis converged on taurodeoxycholic acid (TDCA), a microbiota-derived bile acid that was enriched in human ICT responders in that cohort (31). Together, these results show that antibiotic-induced microbiome changes reshape the host metabolome and identify TDCA as a candidate determinant of ICT responsiveness.

### Pen/Strep treatment significantly depletes TDCA in tumors

Our metabolomics screen identified a bile acid associated with B16-F10 tumor size post ICT (**Fig. 1H**) and with clinical response to ICT therapy in melanoma patients (31). However, the untargeted LC-MS screen detects metabolites based primarily on mass. TDCA and taurochenodeoxycholic acid (TCDCA) are isomeric bile acids with the same molecular mass. As a result, we could not unambiguously distinguish between these two bile acids. TCDCA is a primary bile acid synthesized from cholesterol in the liver, whereas TDCA is a microbiota-derived secondary bile acid (32).

Antibiotics are known to alter bile acid levels by disrupting the gut microbiota and its metabolome (33). In our model, Pen/Strep was the only antibiotic significantly associated with increased tumor size (**Fig. 1C, S1D**), suggesting that the TDCA/TCDCA signal identified in our untargeted analysis was most likely driven by Pen/Strep treatment. Thus, we performed targeted bile acid profiling of tumor and serum samples after Pen/Strep and ICT treatments (**Fig. 2A**). Primary conjugated bile acids tended to increase, while secondary bile acids tended to decrease, in both serum and tumors after Pen/Strep treatment (**Fig. S2B–E, Table S3**). In tumors, TCDCA levels were significantly increased following Pen/Strep treatment (**Fig. 2B, Fig. S2B**), whereas TDCA levels were markedly reduced (**Fig. 2B, Fig. S2C**). Overall, these findings show that Pen/Strep treatment alters the tumor bile acid pool and depletes the microbiota-derived bile acid TDCA.

**Figure 2.**
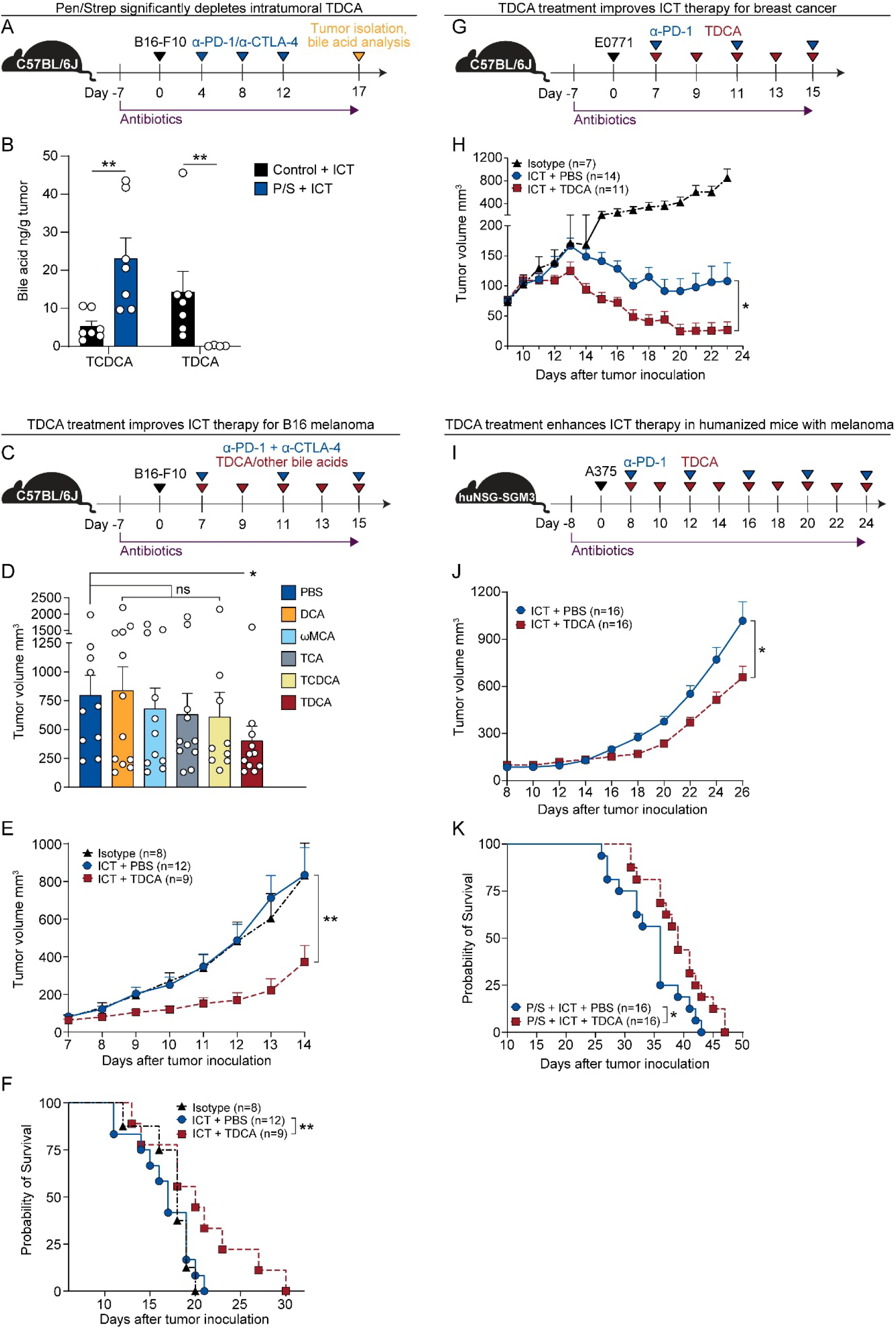
TDCA supplementation overcomes antibiotic-induced ICT hyporesponsiveness across tumor models Schematic overview of tumor bile acid concentration determination. Mice were treated with antibiotics, B16-F10 cells, and ICT as in Figure 1A. Serum and tumors were collected on day 17 post tumor inoculation for targeted bile acid analysis by liquid chromatography-mass spectrometry (LC-MS). Con, no antibiotics control; P/S, Penicillin (0.95 g/L)/ Streptomycin (2 g/L). Concentrations of TCDCA and TDCA in tumors from (A). **(A)** Schematic overview of the B16-F10 heterotopic model with bile acid treatment. Mice (C57BL/6, female, 6–8 weeks old) received penicillin/streptomycin (0.95 g/L; 2 g/L) in drinking water throughout the experiment. 1 x 10^5^ B16-F10 cells were injected subcutaneously into the right inguinal flank. ICT (α-PD-1, α-CTLA-4) or isotype control (rat IgG2a, mouse IgG2b) was injected intraperitoneally, beginning when tumor volumes reached 80 ± 20 mm³. TDCA (50 μl of 4 mM), DCA (50 μl of 4 mM), TCA (50 μl of 16 mM), TCDCA (50 μl of 8 mM), ω-MCA (50 μl of 2 mM) were administrated every two days by intratumoral injection. **(B)** Tumor volumes from day 16 post B16-F10 inoculation, following ICT with treatment of different bile acids (TDCA, DCA, TCA, TCDCA, ω-MCA) as described in (C). *n* =9–11 mice per group. **(C)** Tumor volumes following ICT and antibiotic treatment with or without TDCA from (C); *n* = 8–12 mice per group. **(D)** Survival of tumor-bearing mice from (C); *n* = 8–12 mice per group. **(E)** Schematic overview of the E0771 orthotopic breast cancer model. Mice received penicillin/streptomycin (0.95 g/L; 2 g/L) in drinking water throughout the experiment. 5 x 10^5^ E0771 cells were injected into the mammary fat pad. 200 μg α-PD1 or isotype control were injected intraperitoneally every 4 days, beginning when tumor volumes reached 80 ± 20 mm³. TDCA (50 μl of 4 mM) was administrated every two days by intratumoral injection. **(F)** E0771 tumor volumes from (G); *n* = 7–14 mice per group. **(G)** Schematic overview of the A375 humanized melanoma tumor model. 4–8 weeks old, female, NSG-SGM3 mice were irradiated with 240-250 cGy 24 hours before tail vein transfer of 1 x 10^5^ human hematopoietic CD34^+^ cord blood cells. Mice received penicillin/streptomycin (0.95 g/L; 2 g/L) in drinking water after humanization. 5 x 10^6^ A375 cells were injected subcutaneously into the right inguinal flank. When tumor volumes reached 100 ± 50 mm³, 250 μg human α-PD1 was injected intraperitoneally, every 4 days. TDCA (50 μl of 8 mM) was administrated every two days by intratumoral injection. **(H)** A375 tumor volumes from (I). *n* =16 mice per group. **(I)** Survival of A375 tumor-bearing huNSG-SGM3 mice described in (I); *n* =16 mice per group. DCA, deoxycholic acid. ω-MCA, ω-muricholic acid. TCA, taurocholic acid. TCDCA, taurochenodeoxycholic acid. TDCA, taurodeoxycholic acid. Statistical analysis by Mann-Whitney test (B, J), one-way ANOVA, Kruskal–Wallis test with multiple comparisons (D, E, H), or Log-rank test for (F, K). Points represent individual mice. Bars denote mean ± SEM. All results are representative of ≥2 independent experiments. \**p*< 0.05; \*\****p*** < 0.005; \*\*\****p*** < 0.001; ns, not significant.

### TDCA supplementation overcomes antibiotic-induced ICT hyporesponsiveness across tumor models

Because bile acids can inhibit or enhance antitumor immunity depending on the specific metabolite and/or tumor context (17–22), we asked whether bile acids altered by Pen/Strep treatment modulate antibiotic-induced ICT hyporesponsiveness. Hence, we tested five bile acids whose tumor concentrations were altered by Pen/Strep treatment (**Fig. S2B–C**) and are known to have immunomodulatory effects in cancer (21,23,28,32): two primary conjugated bile acids (TCDCA, taurocholic acid (TCA)) and three secondary bile acids (deoxycholic acid (DCA), ω-muricholic acid (ωMCA), TDCA). We then modified this antibiotic-induced ICT hyporesponsiveness model by administering individual bile acids intratumorally, concurrently with ICT (**Fig. 2C**). Tumor bile acid concentrations increased with bile acid supplementation (**Fig. S3B–F**). Pen/Strep and ICT altered the tumor concentrations of all five bile acids tested (**Fig. S2**), but only TDCA supplementation significantly reduced tumor growth (**Fig. 2D, Fig. S3G**).

We next tested whether TDCA was sufficient to restore ICT responsiveness during antibiotic treatment. Using the same experimental design (**Fig. 2C**), we found that TDCA administration significantly suppressed melanoma growth and prolonged survival compared with vehicle-treated controls (**Fig. 2E–F, Fig. S3H–I**). As expected, Pen/Strep treatment induced ICT hyporesponsiveness, with no significant difference in tumor burden or survival between isotype-and ICT-treated mice (**Fig. 2E–F, Fig. S3H–I**). Importantly, TDCA treatment did not augment ICT when antibiotics were not administered (**Sup. Fig. 3H–I**), clarifying that TDCA only rescues ICT-induced hyporesponsiveness. These results were also consistent with our prior clinical study identifying TDCA as a metabolite associated with improved ICT efficacy in melanoma patients (31).

**Figure 3.**
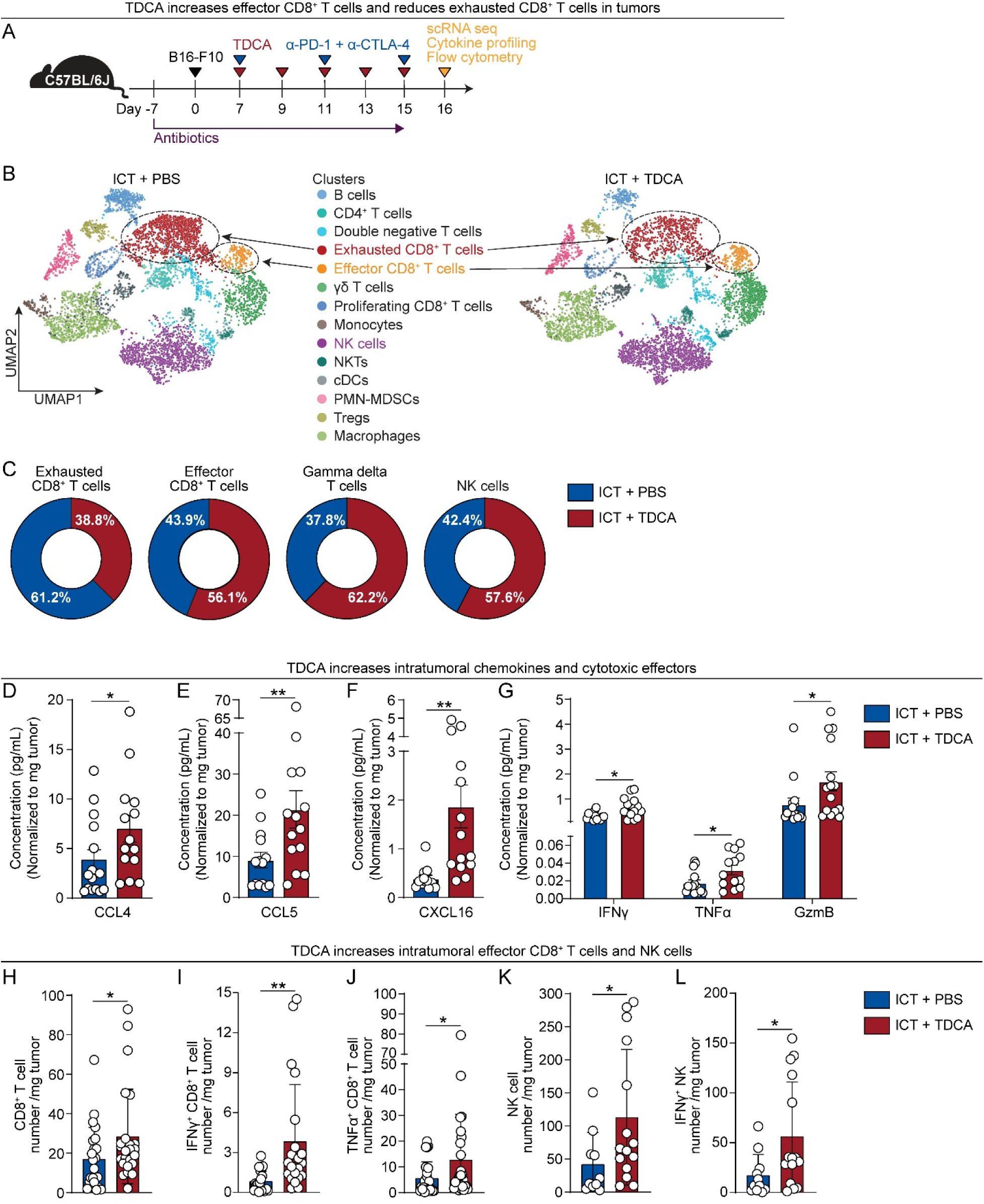
TDCA enhances intratumoral cytotoxic lymphocyte responses during ICT **(A)** Schema of tumor immune profiling experiments. Mice (6–8 week old, female, C57BL6J) received penicillin/ streptomycin (0.95 g/L; 2 g/L) in drinking water throughout the experiment. 1 x 10^5^ B16-F10 cells were injected subcutaneously into the right inguinal flank. ICT was injected intraperitoneally, beginning when tumor volumes reached 80 ± 20 mm³. TDCA (50 μl of 4 mM) was intratumorally injected every two days. Tumors were collected on day 16 post inoculation for immune profiling. **(B)** UMAP plots of immune cell clusters for ICT+PBS and ICT+TDCA treated groups. Clusters were annotated based on canonical gene expression. Dotted circles highlight effector CD8^+^ T cell and exhausted CD8^+^ T cell clusters. **(C)** Proportional contribution of effector CD8^+^ T cell, exhausted CD8^+^ T cell, γδ T cell, and NK cell clusters with or without TDCA administration. **(D-F)** Protein concentrations of chemokines, CCL4 **(D)**, CCL5 **(E)**, CXCL16 **(F)**, and cytotoxic cytokines (IFNγ, TNFα, GzmB) **(G)** from tumor lysates, measured by multiplex Luminex assay. n =13–14 mice per group. **(H-L)** Flow cytometry and quantification of tumor-infiltrating CD8⁺ T cells **(H)**, IFN-γ⁺ CD8⁺ T cells **(I)**, TNF-α⁺ CD8⁺ T cells **(J)**, NK cells **(K)**, and IFN-γ⁺ NK cells **(L)**, normalized to tumor mass. n =10–25 mice per group TDCA, taurodeoxycholic acid; scRNA-seq, single cell RNA sequencing; UMAP, uniform manifold approximation and projection. *Statistical analysis b*y Mann–Whitney test (D-K). Points represent individual mice. Bars denote mean ± SEM. All results (D-K) are representative of ≥2 independent experiments. \**p*< 0.05; \*\***p** < 0.005.

To determine whether the immunomodulatory effects of TDCA extend beyond this melanoma model, we evaluated TDCA supplementation in two additional preclinical tumor models. In an orthotopic E0771 breast cancer model treated with Pen/Strep and α-PD-1, TDCA significantly delayed tumor growth (**Fig. 2G–H**). We next tested TDCA in human melanoma (A375)-bearing humanized NSG-SGM3 mice, an immunologically distinct model containing a functional human immune compartment (**Fig. 2I, Fig. S4**). In this setting, TDCA likewise improved tumor control and prolonged survival during α-PD-1 therapy (**Fig. 2J–K**).

Together, these results demonstrate that TDCA supplementation is sufficient to overcome antibiotic-induced hyporesponsiveness to immune checkpoint therapy across multiple tumor models, identifying TDCA as a key microbiota-derived metabolite that sustains antitumor immunity during ICT.

### TDCA enhances intratumoral cytotoxic lymphocyte responses during ICT

We next sought to determine whether TDCA restrains melanoma tumor growth through a tumor cell-intrinsic mechanism – as DCA, another secondary bile acid, induces apoptosis of colon cancer cells (34). B16-F10 cells were treated with increasing concentrations of TDCA. TDCA had no significant effect on B16-F10 cell growth or cell viability across the doses tested, indicating that its antitumor activity is not tumor cell–intrinsic (**Fig. S5**).

Because TDCA restored ICT responsiveness *in vivo* (**Fig. 2**) without directly affecting tumor cell viability (**Fig. S5**), we next tested whether TDCA enhanced antitumor immunity within the tumor microenvironment. CD45⁺ immune cells were isolated from tumors of ICT-treated mice with or without TDCA treatment and analyzed by single-cell RNA sequencing (scRNA-seq) (**Fig. 3A**). After quality control and lineage annotation (**Fig. 3B, Table S4**), tumors from TDCA-treated mice exhibited increased proportions of effector CD8⁺ T cells, γδ T cells, and NK cells, accompanied by a reduction in exhausted CD8⁺ T cells, compared with vehicle-treated controls (**Fig. 3B–C, Fig. S6B**). These findings suggested that TDCA treatment enhances intratumoral cytotoxic lymphocyte responses, with a particularly prominent effect on CD8^+^ T cell effector function.

To further assess immune activation in the tumor microenvironment, we performed multiplex cytokine and chemokine profiling of tumor lysates. TDCA-treated tumors exhibited elevated levels of lymphocyte-recruiting chemokines CCL4, CCL5 and CXCL16 (35,36) (**Fig. 3D–F**), as well as increased concentrations of effector cytokines IFN-γ, TNF-α, and granzyme B (**Fig. 3G**).

Flow cytometry confirmed that tumors from TDCA-treated mice contained significantly increased numbers of CD8⁺ T cells, including IFN-γ⁺ and TNF-α⁺ CD8⁺ T cells, compared with vehicle-treated controls (**Fig. 3H–J; see Fig. S7A for gating**). TDCA treatment also increased intratumoral NK cells and IFN-γ^+^ NK cells (**Fig. 3K–L**). In contrast, other tumor-infiltrating immune cell populations were not affected by TDCA treatment (**Fig. S6C–K, S7B for gating**). Interestingly, immune cell populations in tumor-draining lymph nodes, including CD8^+^ T cells and NK cells, were unchanged by TDCA treatment (**Fig. S8**). Collectively, these data indicate that TDCA enhances local intratumoral cytotoxic lymphocyte responses during ICT, with the strongest and most consistent effects observed in CD8⁺ T cells.

### TDCA restrains tumor growth by directly enhancing CD8^+^ T-cell cytotoxicity

To determine which lymphocyte populations are required for TDCA-mediated tumor control, we selectively depleted CD4⁺ T cells, CD8⁺ T cells, or NK cells in the B16-F10 melanoma model (**Fig. 4A, S9A–G**). Depletion of CD4⁺ T cells or NK cells did not diminish the antitumor effects of TDCA, whereas CD8⁺ T cell depletion significantly impaired TDCA-mediated tumor control (**Fig. 4B–C, S9H–K**). Notably, CD8⁺ T cell depletion completely abrogated TDCA’s effect, with tumor growth kinetics and survival comparable to vehicle-treated controls (**Fig. 4B–C, Fig. S9H–K**). These findings indicate that, despite the increased intratumoral NK-cell signal observed with TDCA treatment (**Fig. 3K–L**), CD8⁺ T cells are the dominant functional mediators of TDCA-dependent tumor control in this model.

**Figure 4.**
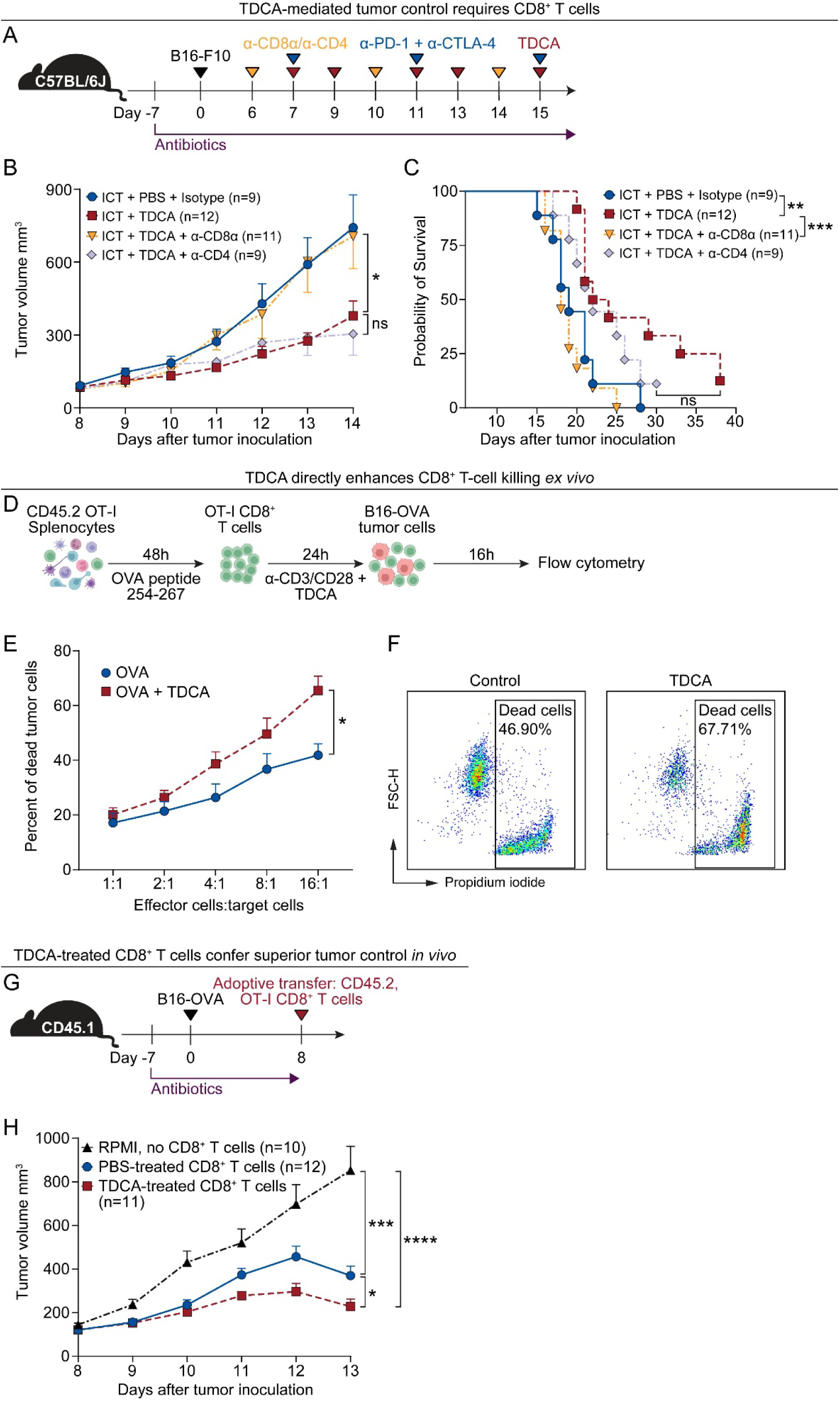
TDCA restrains tumor growth by directly enhancing CD8^+^ T-cell cytotoxicity. **(A)** Schematic overview of CD4^+^/CD8^+^ T cell depletion in B16-F10 tumor model. Mice received penicillin/ streptomycin (0.95 g/L; 2 g/L) in drinking water throughout the experiment. 1 x 10^5^ B16-F10 cells were injected subcutaneously into the right inguinal flank. α-CD4 or α-CD8α (100 μg, intraperitoneal injection) treatments were started one day before ICT treatments. ICT or isotype control treatments began when tumor volumes reached 80 ± 20 mm³. TDCA (50 μl of 4 mM) was intratumorally injected according to the schema. **(B)** Tumor volumes of CD8^+^ T cell– or CD4^+^ T cell–depleted mice from (A); *n* =9–12 mice per group. **(C)** Survival of tumor-bearing, CD8^+^ T cell– or CD4^+^ T cell–depleted mice from (A); *n* =9–12 mice per group. **(D)** Schematic overview of CD8^+^ T cell killing assay. Splenocytes were isolated from CD45.2 OT-I mice and stimulated with OVA peptide 254-267 (10 μg/mL) for 48 hours. CD8^+^ T cells were isolated and stimulated with α-CD3 (2.5 μg/ml), α-CD28 (2.5 ug/ml) and TDCA (200 μM) for 24 hours. Activated CD8^+^ T cells were co-cultured with CFSE-labeled B16-OVA cells for 16 hours. Dead B16-OVA cells were detected by flow cytometry. **(E)** Quantification of the percentage of dead CFSE^+^ B16-OVA cells from (F). Increasing ratios of effector cells (CD8^+^ T cells) to target cells (B16-OVA cells) were tested. **(F)** Representative flow plots of dead B16-OVA from (E). **(G)** Schema of CD8^+^ T cell adoptive transfer. Mice (female, 6–8 weeks old, CD45.2 OT-I) received penicillin/streptomycin (0.95 g/L; 2 g/L) in drinking water throughout the experiment. 2 x 10^5^ B16-OVA cells were injected subcutaneously into the right inguinal flank. CD45.2 OT-I CD8^+^ T cells were activated as described in (D) and 0.5 x10^6^ activated CD8^+^ T cells were transferred by tail vein injection into B16-OVA tumor-bearing CD45.2 OT-I mice when tumor volumes reached 110 ± 30 mm³. **(H)** Tumor volumes following CD8^+^ T cell adoptive transfer from (H); *n* =10–12 mice per group. TDCA, taurodeoxycholic acid. OVA, ovalbumin. Statistical analysis by one-way ANOVA, Kruskal–Wallis test with multiple comparisons (B, H), Mann–Whitney test (E) or Log-rank test (C). Points represent individual mice. Bars denote mean ± SEM. All results are representative of ≥2 independent experiments. \**p*< 0.05; \*\***p** < 0.005; \*\*\***p** < 0.001; ns, not significant.

We next asked whether TDCA directly enhances CD8⁺ T-cell cytotoxicity. To test this in an antigen-restricted system, we used CD8⁺ T cells from OT-I transgenic mice, which recognize a defined ovalbumin (OVA) antigen and thus allow assessment of cytotoxic function without variability in tumor antigen specificity. Activated OT-I CD8⁺ T cells were treated with TDCA and then co-cultured with OVA-expressing B16 melanoma cells (B16-OVA) at varying effector-to-target ratios (**Fig. 4D, Fig. S10**). TDCA-treated CD8⁺ T cells induced significantly greater tumor cell death than control CD8⁺ T cells, indicating enhanced cytotoxic activity (**Fig. 4E–F**). We confirmed this effect *in vivo* using an adoptive transfer model, in which activated TDCA-treated OT-I CD8⁺ T cells were transferred into B16-OVA tumor-bearing mice. Transfer of TDCA-treated CD8⁺ T cells resulted in significantly slower tumor growth than transfer of control CD8⁺ T cells (**Fig. 4G–H**). Together, these results demonstrate that TDCA restrains tumor growth by directly enhancing CD8⁺ T-cell cytotoxic function.

### TDCA enhances CD8^+^ T-cell effector function through bile acid receptor TGR5

The immunological effects of bile acids are mediated through cell surface and nuclear receptors. Thus, we next sought to define the receptor through which TDCA directly enhances CD8⁺ T-cell effector function. We tested three bile acid receptors (using luciferase reporter assays) with established immunologic relevance— Takeda G protein-coupled receptor 5 (TGR5) (37–40), farnesoid X receptor (FXR) (25,26), and vitamin D receptor (VDR) (25,27,41,42)—across a range of TDCA doses. TDCA strongly activated TGR5 at low doses, induced only modest activation of FXR, and did not activate VDR (**Fig. 5A–C**), identifying TGR5 as a principal receptor mediating TDCA signaling.

**Figure 5.**
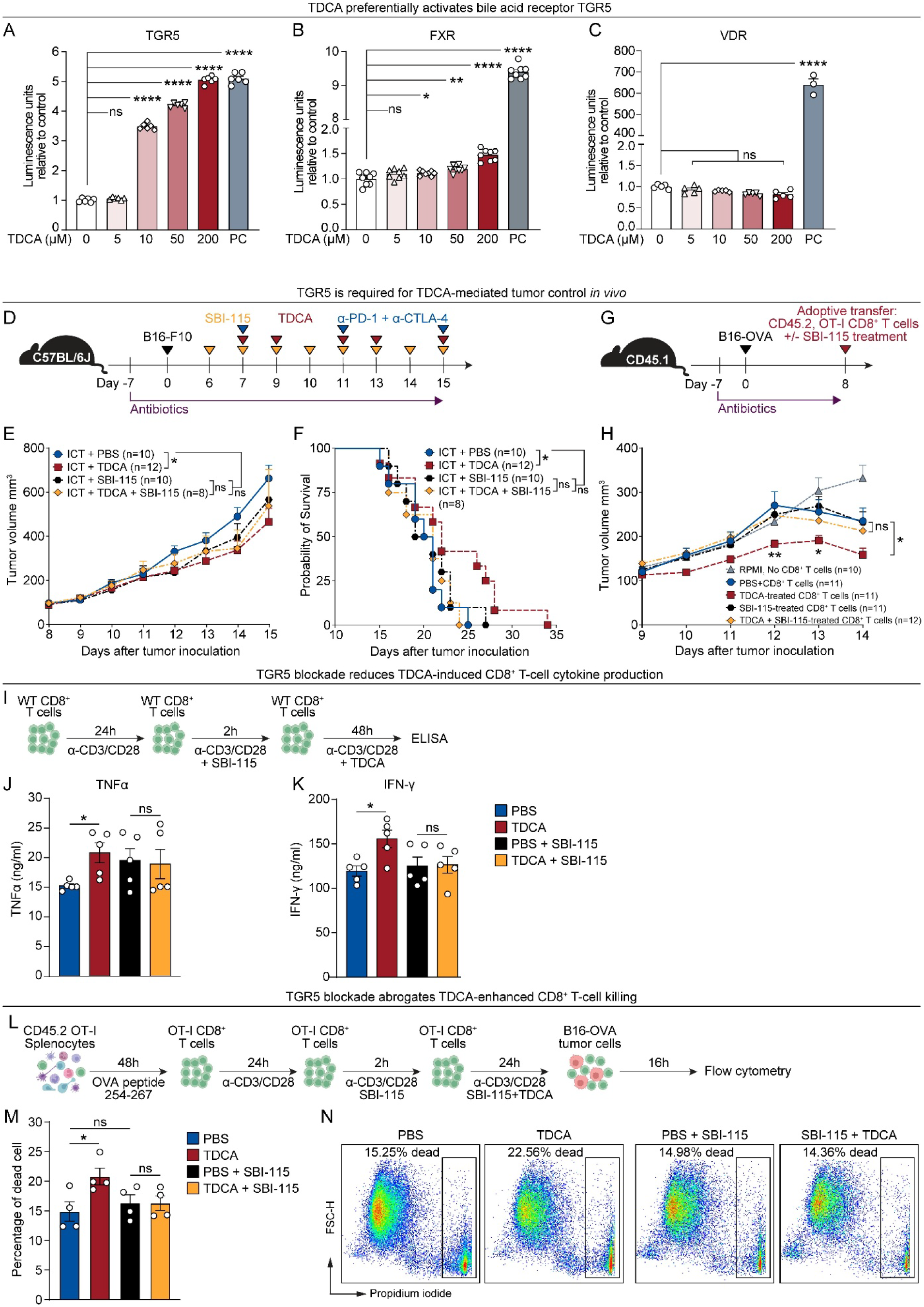
TDCA enhances CD8^+^ T-cell effector function through bile acid receptor TGR5 (A-C) Dose-dependent activation of bile acid receptors by TDCA in luciferase reporter assays. Reporter activity was normalized to 0 μM TDCA. **(A)** HEK293T cells were transfected with a TGR5 luciferase reporter. 10 μM of proprietary Cayman Chemical TGR5 agonist was used as a positive control. **(B)** HEK293T cells were transfected with a FXR luciferase reporter. 1 μM XL335 was used as a positive control. **(C)** Proprietary engineered human cells with high expression of the human VDR and a luciferase reporter were tested for TDCA activation; 0.25 mM calcitriol as a positive control. **(D)** Schema of the effect of TGR5 antagonist SBI-115 on B16-F10 tumor model. Mice (6–8 weeks old, female, C57BL/6) received penicillin/streptomycin (0.95 g/L; 2 g/L) in drinking water throughout the experiment. 1 x 10^5^ B16-F10 cells were injected subcutaneously into the right inguinal flank. ICT or isotype control treatments began when tumor volumes reached 80 ± 20 mm³. TDCA (50 μl of 4 mM) was intratumorally injected according to the schema. TGR5 antagonist SBI-115 (10 mg/kg) was intraperitoneally injected once a day, starting one day before TDCA treatments. **(E)** Tumor volumes from treated mice in (D); *n* =8–12 mice per group. **(F)** Survival of tumor-bearing mice from (D); *n* =8–12 mice per group. **(G)** Schematic overview of SBI-115 and TDCA-treated CD8^+^ T cell adoptive transfer. Splenocytes were isolated from CD45.2 OT-I mice and stimulated with OVA peptide 254-267 (10 μg/mL) for 48 hours. CD8^+^ T cells were isolated and stimulated with α-CD3 (2.5 μg/ml) and α-CD28 (2.5 ug/ml) for 24 hours. The activated CD8^+^ T cells were then treated with TGR5 antagonist SBI-115 (10 μg/ml), for 2 hours before TDCA treatment (200 μM) for 24 hours. α-CD3 (2.5 μg/ml) and α-CD28 (2.5 ug/ml) were included the treatment media for SBI-115 and TDCA. 0.5 x 10^6^ SBI-115 and TDCA-treated CD45.2 OT-I CD8^+^ T cells were transferred by tail vein injection into B16-OVA tumor-bearing CD45.2 OT-I mice (females, 6–8 weeks old) when tumor volumes reached 110 ± 30 mm³. **(H)** Tumor volumes from B16-OVA tumor-bearing mice from (G); *n* =10–12 mice per group. **(I)** Schema of CD8^+^ T cell preparation for TDCA-induced T-cell cytokine secretion. CD8^+^ T cells were isolated from C57BL/6 mice (females, 6-8 weeks old) and stimulated with α-CD3 (2.5 μg/ml) and α-CD28 (2.5 μg/ml) for 24 hours. The activated CD8^+^ T cells were then treated with TGR5 antagonist SBI-115 (10 μg/ml) for 2 hours before TDCA treatment (200 μM) for 48 hours (α-CD3 and α-CD28 included in the treatment media). Supernatants were collected for cytokine detection by ELISA. **(J-K)** ELISAs for cytotoxic cytokines with supernatants from TGR5 antagonist-and TDCA-treated CD8^+^ T cell cultures. **(J)** TNF-α and **(K)** IFN-γ **(J)** Schematic overview of T cell killing assay with TGR5 blockade (SBI-115 treatment). Splenocytes were isolated from CD45.2 OT-1 mice and stimulated with OVA peptide 254-267 (10 μg/mL) for 48 hours. Activated CD8^+^ T cells were isolated and stimulated with α-CD3 (2.5 μg/ml) and α-CD28 (2.5 ug/ml) for 24 hours. The activated CD8^+^ T cells were then treated with TGR5 antagonist SBI-115 (10 μg/ml) for 2 hours before TDCA treatment (200 μM) for 24 hours (α-CD3 and α-CD28 included in the treatment media). Activated CD8^+^ T cells were co-cultured with CFSE-labeled B16-OVA cells for 16 hours. Dead B16-OVA cells were detected by flow cytometry. **(K)** Quantification of the percentage of dead CFSE^+^ B16-OVA cells from (L). **(L)** Representative flow plots of dead B16-OVA cells from (L). TGR5, Takeda G protein-coupled receptor 5; FXR, farnesoid X receptor; VDR, vitamin D receptor; TDCA, taurodeoxycholic acid. Statistical analysis by ordinary one-way ANOVA (A-C), one-way ANOVA, Kruskal–Wallis test (E, H, J, K, M) or Log-rank test for (F). Points represent individual mice. Bars denote mean ± SEM. All results are representative of ≥2 independent experiments. \**p*< 0.05; \*\***p** < 0.005; \*\*\***p** < 0.001; ns, not significant.

To test whether TGR5 signaling is required for TDCA-mediated tumor control *in vivo*, we inhibited TGR5 with the antagonist SBI-115 during ICT and antibiotic treatments (**Fig. 5D**). Pharmacological antagonism of TGR5 with SBI-115 mitigated TDCA-mediated tumor control and survival benefit (**Fig. 5E–F**). To determine whether this effect was CD8^+^ T cell intrinsic, we pre-treated activated OT-I CD8⁺ T cells with SBI-115 before TDCA exposure and then transferred the cells to B16-OVA tumor-bearing mice (**Fig. 5G**). TGR5 antagonism blunted the antitumor activity of TDCA-treated CD8^+^ T cells (**Fig. 5H**). Together, these results suggest that TDCA requires TGR5 signaling to enhance CD8^+^ T cell-mediated tumor control *in vivo*.

We next asked whether TGR5 directly mediates TDCA-induced CD8⁺ T cell effector function. TGR5 antagonism with SBI-115 reduced TDCA-induced IFN-γ and TNF-α production by CD8^+^ T cells (**Fig. 5I–K**). In the *ex vivo* OT-I T cell killing assay, pre-treatment with SBI-115 before TDCA exposure (**Fig. 5L**) also decreased TDCA-mediated CD8⁺ T cell tumor killing, compared to the vehicle control (**Fig. 5M–N**). These results demonstrate that TDCA directly augments CD8⁺ T cell effector and cytotoxic function through the bile acid receptor TGR5.

## Discussion

In this study, we identify the microbiota-derived bile acid taurodeoxycholic acid (TDCA) as a key metabolite that restores ICT efficacy against melanoma after antibiotic-induced hyporesponsiveness. Using antibiotic perturbation coupled with metabolomic profiling, we found that distinct antibiotics generated different metabolomic states and heterogeneous tumor phenotypes during ICT. Penicillin/streptomycin (Pen/Strep), the antibiotic regimen most strongly associated with impaired tumor control, significantly reduced intratumoral TDCA. TDCA supplementation alone was sufficient to overcome antibiotic-induced ICT hyporesponsiveness across multiple tumor models. Mechanistically, TDCA increased tumor-infiltrating effector CD8⁺ T cells, enhanced effector cytokine production, and directly augmented CD8⁺ T-cell cytotoxicity through the bile acid receptor TGR5. Together, these findings identify a microbiota-derived bile acid–TGR5–CD8⁺ T-cell axis as a determinant of immunotherapy responsiveness.

The gut microbiota is now recognized as an important regulator of ICT response in both preclinical models and patients (5–15,31). Antibiotic exposure has been repeatedly associated with impaired immunotherapy responses across multiple cancer types (13–16). Our data extend these observations by showing that antibiotic effects are not uniform: distinct antibiotic regimens produced different degrees of microbiota disruption, divergent metabolomic consequences, and heterogeneous tumor phenotypes during ICT. These findings provide a plausible mechanistic explanation for why the clinical impact of antibiotics on immunotherapy has been variable across studies (43). More broadly, our results support a model in which microbiota-dependent metabolites are not merely byproducts of commensal communities, but active mediators that help sustain antitumor immunity.

Microbiota-derived metabolites(44–50)—including inosine (44,50) and short-chain fatty acids (45,46)—enhance antitumor immunity through a variety of mechanisms. Inosine produced by the commensal *Bifidobacterium psuedolongum*, together with a costimulus, enhances ICT efficacy by promoting T_H_1 differentiation and effector functions (44). Similarly, commensal-derived short-chain fatty acids pentanoate and butyrate induce a glycolytic switch in CD8^+^ T cells, improving cytotoxic cytokine secretion, *in vivo* survival and proliferation, and tumor control (46). Butyrate treatment combined with chemotherapy improves control of colon cancer through the ID2–IL-12 axis in CD8^+^ T cells (47). *Lactobacillus reuteri*, a common probiotic species, translocates to tumors and produces indole-3-aldehyde, which activates the transcription factor AhR in CD8^+^ T cells to drive cytotoxic gene expression and antitumor immunity. Indole-3-aldehyde treatment enhances α-PD-L1 efficacy in a preclinical melanoma model and is enriched in responding melanoma patients, highlighting the importance of microbial metabolites in cancer immunotherapy (48). Collectively, these microbial metabolites converge on T-cell differentiation, activation, or effector function modulation. Our findings place TDCA within this emerging framework, but with an important distinction: rather than broadly associating a metabolite with a tumor response, we show that TDCA is sufficient to rescue antibiotic-induced ICT hyporesponsiveness and directly enhance CD8⁺ T-cell cytotoxicity. These findings position TDCA as a defined and mechanistically tractable mediator of microbiota-dependent immunotherapy response.

Direct links between bile acid signaling and antitumor immunity have recently begun to emerge and suggest that bile acid effects are highly context dependent. For example, the microbiota-derived secondary bile acid deoxycholic acid (DCA) inhibits Ca²⁺-NFAT signaling in CD8⁺ T cells, suppressing effector functions and promoting colorectal cancer growth (21). In a SV40 large T antigen (TAG) tumorigenic liver cancer model, elevated bile acid synthesis in the tumor microenvironment leads to accumulation of conjugated bile acids, including TCDCA and LCA. The high concentrations of TCDCA induce reactive oxygen species in CD8⁺ T cells, leading to T cell death and loss of α-PD-1 efficacy. Similarly, LCA conjugates induce endoplasmic reticulum stress in CD8^+^ T cells, causing reduced tumor control (23). While secondary bile acids are immunosuppressive in B16 liver tumors, primary bile acids increase hepatic endothelial expression of CXCL16 and infiltration of CXCR6^+^ natural killer T (NKT) cells, thus promoting tumor control (32). Similarly, dietary supplementation with UDCA, a secondary unconjugated bile acid, was found to reduce liver tumor size (23). Last, microbiota-conjugated secondary bile acid isomers antagonize the androgen receptor in CD8⁺ T cells, promoting a stem-like phenotype and enhancing ICT efficacy against bladder cancer (28). Our work adds to this evolving literature by identifying TDCA as a bile acid that supports antitumor immunity, and by linking its activity specifically to the rescue of antibiotic-induced ICT hyporesponsiveness in melanoma. These differences across studies likely reflect important effects of bile acid identity, dose, receptor usage, and tumor context.

While bile acids are increasingly recognized as potent immunomodulatory signaling molecules (17–23,51), few studies have examined T cell immunomodulation by TDCA. Notably, *in vitro* TDCA treatment of activated T cells specific for TAG (an oncoprotein) failed to increase IFNγ^+^TNFα^+^ T cell numbers (23). This contrasts with our findings that TDCA increases IFNγ^+^ and TNFα^+^ secretion and tumor killing by CD8^+^ T cells. These alternate results likely reflect differences in TDCA doses (200 μM vs 100 μM), antigen models (OVA vs TAG (SV40 large T cell antigen)), and readouts (cytotoxicity and cytokine production vs T cell numbers). Further, several studies identified an indirect, immunosuppressive role for TDCA through myeloid cells. In an obesity mouse model, TDCA and valine administration suppressed macrophage inflammation via a TGR5-dependent mechanism, leading to decreased T-cell activation (51). Similarly, TDCA treatment during DSS colitis reduced the number of inflammatory macrophages and T cells in the lamina propria and inhibited *in vitro* inflammasome activation in macrophages (52). TDCA increased the number of myeloid-derived suppressor cells in a sepsis model, improving survival of mice (53). These examples demonstrate that TDCA suppresses inflammation in myeloid cells and likely has context-dependent activation of T cells.

Mechanistically, we found that TDCA enhances CD8⁺ T-cell effector function through TGR5. Pharmacologic inhibition of TGR5 mitigated TDCA-mediated rescue of tumor control *in vivo* and reduced TDCA-induced cytokine production and tumor killing by CD8⁺ T cells *ex vivo*. The downstream signaling pathways remain to be defined. TGR5 is a G-coupled receptor known to induce accumulation of cAMP and activation of Protein Kinase A (PKA) (54). However, the effects of the TGR5-cAMP pathway on CD8⁺ T-cell cytotoxicity are highly context-dependent. Prior work has implicated TGR5 signaling in both immunosuppressive myeloid programs and pro-effector CD8⁺ T-cell responses in other disease settings (51–56). Based on this, we speculate that TDCA may induce the TGR5-cAMP-mTOR pathway to promote cytotoxic CD8^+^ T cells and antitumor immunity in our melanoma model. Future studies will be needed to define the downstream transcriptional, immune, and metabolic programs involved.

This study has several limitations. First, the mechanistic work was performed predominantly in preclinical mouse models, and the human relevance of TDCA is supported by prior correlative clinical data rather than by prospective patient validation. Second, although antibiotic treatment depletes TDCA and supplementation rescues response, the specific microbial taxa and enzymatic pathways driving this loss remain undefined. Third, our mechanistic studies relied on pharmacologic antagonism of TGR5 rather than genetic deletion and do not fully exclude contributions from TGR5 signaling in additional cell types or from off-target effects. Finally, the activity of TDCA may vary by tumor type, immune context, and bile acid dose. These questions will be important to address in future translational and mechanistic studies.

Our study helps to distinguish microbiota-derived metabolites from live-bacteria–based approaches to improving immunotherapy response. Fecal microbiota transplantation and supplementation with beneficial bacterial strains remain important experimental and clinical strategies (5,8,12), but they introduce challenges related to engraftment, reproducibility, safety, and regulatory complexity. By contrast, our findings suggest that replacement of defined microbial metabolites may provide a more tractable approach to restoring beneficial microbiota functions after antibiotic exposure. In that sense, TDCA may represent a proof-of-principle for metabolite replacement as a strategy to mitigate antibiotic-associated loss of immunotherapy response.

In summary, our study identifies TDCA as a microbiota-derived metabolite that directly enhances CD8⁺ T-cell–mediated anti-tumor immunity and restores responsiveness to immune checkpoint therapy following antibiotic-induced dysbiosis. These findings provide mechanistic insight into how microbial metabolic products regulate cancer immunotherapy outcomes and highlight bile acid signaling as a potential therapeutic axis for enhancing immunotherapy efficacy.

More broadly, our work supports the concept that targeting microbiota-derived metabolites represents a tractable strategy to enhance immunotherapy responses in patients with microbiota disruption.

## Acknowledgments

We thank Dr. Yangxin Fu for providing B16-OVA cells. We thank Indu Raman from the UT Southwestern Microarray Core for assistance with Luminex assays. We thank Mauricio Velasquez from the UT Southwestern Lipid Mass Spectrometry Core for the targeted bile acid analysis. We thank the UT Southwestern McDermott Next Generation Sequencing Core for assistance with scRNA-sequencing. Components of some figures were made using BioRender.

## Author contributions

W.L. and A.Y.K. designed the research. W.L., H.W., P.S., P.D.V., L.A.C., N.P., J.N.L., R.S.-Y.C., T.S., C.Z., M.J.V., I.R., J.M., V.J.M., R.K., P.R., and X.V.L. performed the research. W.L., S.G., C.L., J.K., X.Z. and J.B.X. analyzed the data. W.L., C.M.Z., and A.Y.K. wrote the paper. C.M.Z. and A.Y.K. edited the paper. All authors revised the manuscript and approved its final version.

## Data and materials availability

Single cell RNA sequencing are available from the Gene Expression Omnibus (GEO) repository under GSE327301. Metagenomic shotgun sequencing data are deposited under PRJNA1450123 in the NCBI Sequence Read Archive (SRA): http://ncbi.nlm.nih.gov/sra. Metabolomics shotgun and targeted analyses will be available from the National Metabolomics Data Repository (NMDR) (data deposition ongoing, please refer to **Tables S1-3**). All other data are available in the main text or the supplementary materials.

## Code Availability

Code for MSS, scRNA seq, and untargeted metabolomics analyses can be accessed at https://github.com/zhanxw/tdca-paper-code.

## Materials and Methods

### Cell Lines

B16-F10 cells (ATCC CRL-6475; RRID:CVC_0159), B16-OVA cells (gift from Dr. Yangxin Fu, School of Basic Medical Sciences, Tsinghua University, RRID:CVCL_WM78), A375 cells (ATCC CRL-1619, RRID:CVCL_0132) and EO771 cells (ATCC CRL-3461; RRID:CVCL_GR23), were grown at 37°C under 5% CO2 in DMEM medium (Gibco, #11995-065) supplemented with 10% heat-inactivated FBS (Corning, #MT35016CV). All cell lines routinely tested negative for *Mycoplasma* (ATCC PCR Mycoplasma Detection Kit, #30-1012K). Only cells with a passage number <10 were used for experiments.

### Mice

C57BL/6J (Stock No: 000664), CD45.1 C57BL/6 (Stock No: 002014), C57BL/6-Tg (TcraTcrb)1100Mjb/J (OT-1, Stock No: 003831) and NSG-SGM3 (Stock No: 013062) mice were obtained from Jackson Laboratories and bred and maintained in the barrier facility at the University of Texas Southwestern Medical Center. All animals were kept on a 12-hour light-dark cycle and were fed standard mouse chow (Teklad 2916, irradiated). Sex-matched, 6-8 week old mice were used for all experiments and co-housed littermates were used as controls. Experiments were performed using protocols approved by the Institutional Animal Care and Use Committees of the UT Southwestern Medical Center.

### Humanized NSG-SGM3 (huNSG-SGM3) mouse model

The humanized NSG-SGM3 model was generated as previously reported (57). Briefly, NSG-SGM3 mice (female, 4-8 weeks old, Jackson #013062) were irradiated with 240-250 cGy 24 hours before tail vein transfer of 1x 10^5^ human hematopoietic CD34^+^ cord blood cells. CD34^+^ cord blood cells were acquired from Lonza (Cat #2C-101 or 2C-101H). 12 weeks after human CD34^+^ cell transfer, the engraftment of the human immune system was evaluated by flow cytometry of peripheral blood. A mouse was considered “humanized” if greater than 25% of the total CD45^+^ cells in peripheral blood express human CD45.

### Preclinical tumor mouse models

#### Tumor monitoring and endpoints for all models

Tumor volumes were measured every 1-2 days using a digital caliper. Tumor volumes were calculated using the formula: (length x width x width)/2. Loss of survival was defined as death (with moribund mice being euthanized), when tumor diameter exceeded 2 cm in any dimension, or when the tumor volumes exceeded 1,800 mm^3^.

#### Metabolite screening assay with melanoma model

Female C57BL/6J mice (6-8 weeks old) provided one of six antibiotic treatments, starting one week before tumor inoculation and continuing throughout the experiment. Penicillin G (0.95 g/L, Sigma-Aldrich, #P8721-100MU)/streptomycin sulfate (2 g/L, Thermo Fisher Scientific, #AC455341000), vancomycin hydrochloride (0.5 g/L, Thermo Scientific Chemicals, #AAJ6279006), levofloxacin (0.5 g/L, Fagron, NDC #51552-1503-3), and clindamycin (0.5 g/L, Research Products International, #50213243) were administrated to mice via drinking water, *ad libitum*. Piperacillin/tazobactam (12.5 mg/ml, Auromedics, NDC #55150-121-50) was intraperitoneally injected once a day and cefepime (12.5 mg/ml, Apotex Corp, NDC #60505-6146-4) was intraperitoneally injected twice a day. 1 x 10^5^ B16-F10 cells were implanted subcutaneously into the right flank of mice on day zero. 200 µg of α–PD-1 (clone RMP1-14, Bio X Cell, #BE0146) and 200 µg α–CTLA4 (9D9, Bio X Cell, #BP0164) or isotype control (rat IgG2a, Bio X Cell, #BE0089; mouse IgG2b, Bio X Cell, #BP0086) antibodies were intraperitoneally injected on days 4, 8 and 12.

#### Preclinical melanoma model with bile acid treatment

Female C57BL/6J mice (6-8 weeks old) were given penicillin/streptomycin (0.95 g/L; 2 g/L) in drinking water, *ad libitum*. Pen/strep was provided for the duration of the experiment. Seven days after antibiotic treatment was started, 1 ×10^5^ B16-F10 cells were implanted subcutaneously into the right flank of mice. On day 7 after tumor inoculation, mice were separated into treatment groups with equivalent tumor volumes (80mm^3^ ± 20 mm^3^). Mice that fell outside the tumor size criteria were excluded from the experiment. Following group assignment on day 7, mice were intraperitoneally injected with 200 μg anti-PD-1 (RMP1-14, Bio X Cell, #BE0146) and 200 μg anti-CTLA-4 (9D9, Bio X Cell, #BP0164) antibodies or 200 μg isotype control (rat IgG2a, Bio X Cell, #BE0089; mouse IgG2b, Bio X Cell, #BP0086, respectively) every four days (days 7, 11, 15). Bile acids TDCA (50 μl of 4 mM, Sigma, #580221), DCA (50 μl of 4 mM, Sigma, #D2510), TCA (50 μl of 16 mM, Sigma, #580217), TCDCA (50 μl of 8 mM, Sigma, #T6260), or ω-MCA (50 μl of 2 mM, Steraloids, #C1888-000) were administrated every two days by intratumoral injection, starting on day 7 post tumor inoculation. PBS (Gibco, #10010-049) was used as a vehicle control for bile acid treatment. Tumors were collected on day 16 post inoculation for volume measurements and bile acid analysis.

#### E0771 breast cancer preclinical model

Female C57BL/6J mice (6-8 weeks old) were given penicillin/streptomycin (0.95 g/L; 2 g/L) in drinking water, *ad libitum*. Pen/strep was provided for the duration of the experiment. Seven days after antibiotic treatment was started, 5 x 10^5^ E0771 cells were injected into the mammary fat pad. 200 μg α-PD1(RMP1-14, Bio X Cell, #BE0146) or isotype control (Rat IgG2a, Bio X Cell, #BE0089) antibodies, were injected intraperitoneally every 4 days, beginning when tumor volumes reached 80 ± 30 mm³. TDCA (50 μl of 4 mM, Sigma, #580221) was administrated every two days by intratumoral injection.

#### Human A375 melanoma in humanized NSG-SGM3 (huNSG-SGM3) mice model

huNSG-SGM3 mice were generated and validated. At 16-20 weeks old (12 weeks post human stem cell transfer), mice were given penicillin/streptomycin (0.95 g/L; 2 g/L) in drinking water, *ad libitum*. Pen/Strep was provided for the duration of the experiment. Eight days after antibiotic treatment was started, 5 x 10^6^ A375 cells were injected subcutaneously into the right inguinal flank. When tumor volumes reached 100 ± 50 mm³, 250 μg human α-PD-1 (RMP1-14, Bio X Cell) was injected intraperitoneally every 4 days. TDCA (50 μl of 8 mM, Sigma, #580221) was administrated every two days by intratumoral injection. PBS was used as a vehicle control for TDCA treatment. Tumor volumes and survival were monitored for up to 48 days post tumor injection.

#### Immune cell depletion assays

The preclinical melanoma model with TDCA treatment was used. Starting day 6 post tumor inoculation, 100 μg of α-CD8α (clone 2.43, Bio X Cell, #BE0061), 100 μg α-CD4 (clone GK1.5, Bio X Cell, #BE0003-1), 250 μg of α-NK1.1 (clone PK136, Bio X Cell, #BE0036) or isotype control (Rat IgG2b, Bio X Cell, #BE0090 for α-CD8α and α-CD4; Mouse IgG2a, Bio X Cell, #BE0085 for α-NK1.1) were injected intraperitoneally every 4 days.

#### TGR5 blockade with SBI-115

The preclinical melanoma model with TDCA treatment was used. TGR5 antagonist SBI-115 (10 mg/kg, MedChemExpress, #HY-111534) was intraperitoneally injected once a day, starting six days post tumor inoculation (one day before TDCA treatments).

### *In vitro* T cell effector cytokine detection

CD8^+^ T cells were isolated from spleens of C57BL/6J mice (8-12 weeks old) using the mouse EasySep CD8^+^ T cell isolation kit (STEMCELL Technologies, #19853). 1 x 10^6^ T cells were cultured in RPMI-1640 (Gibco, #11875-119) supplemented with 10% FBS (Corning, #MT35015CV), 1% pencillin/streptomycin (10,000 U/mL, Gibco, #15140-122), 10 mM HEPES (Gibco, #15630-080), 1 mM sodium pyruvate (Gibco, #11360-070), 50 μM 2-Mercaptoethanol (Gibco, #31350010) and 1% MEM non-essential amino acids (Gibco, #11140050). The cells were activated overnight with α-CD3ε (2.5 μg/ml, clone145-2C11, Bio X Cell, #BE0001-1) and α-CD28 (2.5 μg/ml, clone PV-1, Bio X Cell, #BE0015-5) antibodies. α-CD3ε (2.5 μg/ml) and α-CD28 (2.5 μg/ml) were included in all following treatment medias. Activated CD8^+^ T cells were treated with TGR5 antagonist SBI-115 (10 ng/ml, MedChemExpress, #HY-111534) for 2 hours and then treated with TDCA (20 μM) for 48 hours. The supernatant was collected and IFN-γ and TNFα expression was determined by ELISA (Mouse IFN-gamma DuoSet ELISA, R&D Systems, #DY485-05; Mouse TNF-alpha DuoSet ELISA, R&D Systems, #DY410-05).

#### T cell killing assays

Splenocytes were isolated from OT-I mice and cultured in T cell culture medium (RPMI-1640 (Gibco, #11875-119) supplemented with 10% FBS (Corning, #MT35015CV), 1% pencillin/streptomycin (10,000 U/mL, Gibco, #15140-122), 10mM HEPES (Gibco, #15630-080), 1mM sodium pyruvate (Gibco, #11360-070), 50 μM 2-mercaptoethanol (Gibco, #31350010) and 1% MEM non-essential amino acids (Gibco, #11140050)) with OVA SIINFEKL peptide (10 μg/ml, Sigma-Aldrich, #S7951) for 48 hours. CD8^+^ T cells were isolated using the mouse EasySep CD8^+^ T cell isolation kit (STEMCELL Technologies, #19853) and treated with TDCA (200 μM), α-CD3ε (2.5 μg/ml) and α-CD28 (2.5 μg/ml) for 24 hours. B16-OVA cells were labeled with 0.25 μM CFSE (Biolegend, #423801) and plated. Three hours later, TDCA-treated CD8^+^ T cells were added to the CFSE-labeled B16-F10 cells at 1:1, 2:1, 4:1, 8:1, and16:1 (effector CD8^+^ T cell: B16-OVA target cell) ratios. 16 hours later, viability of B16-OVA was determined by flow cytometry.

For TGR5 blockade by antagonist SBI-115, CD8^+^ T cells were isolated from splenocytes and stimulated with OVA SIINFEKL peptide as described above, with a few exceptions. After OVA stimulation, CD8^+^ T cells were treated with α-CD3ε (2.5 μg/ml, 145-2C11, Bio X Cell, #BE0001-1) and α-CD28 (2.5 μg/ml, PV1, Bio X Cell, #BE0015-5) for 24 hours. SBI-115 (10ng/ml, MedChemExpress) was added to the culture for 2 hours and then CD8^+^ T cells were treated with TDCA (20 μM), α-CD3ε (2.5 μg/ml) and α-CD28 (2.5 μg/ml) for 24 hours. CFSE labeled-B16-F10 were co-cultured with treated, activated CD8^+^ T cells at 1:1 ratio for 16 hours. Viability of B16-OVA cells was detected by flow cytometry.

#### Adoptive T cell transfer

CD45.1 C57BL/6 mice were provided penicillin/streptomycin (0.95 g/L; 2 g/L) in drinking water, *ad libitum*, throughout the experiment, starting seven days before tumor inoculation. 2 x 10^5^ B16-OVA tumor cells were subcutaneously injected into the right flank of the mice. On day 8 after tumor inoculation, mice were separated into treatment groups with equivalent tumor volumes (110mm^3^ ± 40mm^3^). Mice not meeting tumor size criteria were excluded from the experiment. CD8^+^ T cells were activated and treated as described for the *in vitro* T cell killing assays. 0.5 × 10^6^ OT-I CD8^+^ T cells with or without TDCA (200 μM) and SBI-115 (10 ng/ml) treatment were transferred into the tumor-bearing CD45.1 mice. Tumor volumes were monitored throughout the experiment.

#### MTS assay

B16-F10 cells were seeded in 96 well plates at a concentration of 1 x 10^3^ cells/well with different doses of TDCA (0 μM, 100 μM, 200 μM, 400 μM). Cell proliferation was monitored for 5 days with MTS reagent (Abcam, #ab197010). MTS reagent was added every day and the absorbance at 490 nm was measured every 30 min for 4 h at 37°C on a Biotek Synergy LX (Agilent). Absorbance readings were normalized to day 1 absorbance.

### Luciferase cell reporter assays

#### TGR5 luciferase reporter

TDCA-induced TGR5 activation was measured using the TGR5 Reporter Assay Kit (Cayman Chemical, #601440), following the manufacturer’s instructions. HEK293T cells were seeded on the assay plates and treated with TDCA (0 μM, 10 μM, 50 μM, 200 μM) or 10 μM TGR5 agonist (Cayman Chemical) for 16-18 hours. Luminescence was measured using a microplate reader (Biotek Synergy LX, Agilent).

#### FXR luciferase reporter

TDCA-induced FXR activation was measured using the FXR Reporter Assay Kit (Cayman Chemical, #601790), following the manufacturer’s instructions. HEK293T cells were seeded on the assay plates and treated with TDCA (0 μM, 10 μM, 50 μM, 200 μM) or 1 μM XL335 (FXR positive control, Cayman Chemical) for 16-18 hours. Luminescence was measured using a microplate reader (Biotek Synergy LX, Agilent).

#### VDR luciferase reporter

TDCA-induced VDR activation was measured using a human VDR (NR1I1) reporter assay (INDIGO Biosciences, #IB00701), following manufacturer’s instructions for an agonist-mode assay. Briefly, proprietary reporter cells were treated for 22–24 hours with TDCA (0 μM, 10 μM, 50 μM, 200 μM) or calcitriol (0.25 nM, INDIGO Biosciences) as a positive control. Luciferase activity was measured using a plate reader (Biotek Synergy LX, Agilent). Cell-free wells containing medium only were used to quantify background luminescence.

### Metabolome extraction for Liquid Chromatography-Mass Spectrometry analysis

#### Serum metabolites

Whole blood was collected in serum separator tubes (BD, #367983) and allowed to clot for 30 min at room temperature. Samples were centrifuged at 2,000 x g for 10 min at 4°C, and serum was collected. 20 µL of serum was mixed with 180 µL of 80% methanol (Optima LC-MS grade, Fisher Chemical, #A456-500), incubated at 4°C for 1 h, and centrifuged at 14,000 x g for 15 min. The clarified supernatants were transferred to fresh tubes and stored at-80°C until LC-MS analysis.

#### Fecal metabolites

Fecal metabolites were extracted using an organic solvent–based protocol. Briefly, 40–50 mg of mouse feces was placed into microcentrifuge tubes, ensuring that sample weights across the cohort were within a 1–1.5-fold range. Metabolites were extracted using a 2:1 (v/v) acetone:isopropanol extraction solvent (Suprasolv Acetone for GC MS,Supelco, #1.00658.1000; LiChrosolv 2-Propanol gradient garde for LC, Supelco, #1.01040.1000). For each sample, 300 μL of room-temperature extraction solvent was added and samples were vortexed for 15 seconds twice. The supernatant was transferred to fresh microtubes and temporarily stored at-80°C while the remaining samples were processed. The extraction was repeated once more with an additional 300 μL solvent, and the second supernatant was combined with the first extract, yielding approximately 600 μL total extract per sample. The pooled extracts were centrifuged at 13,500 x g for 10 minutes to pellet insoluble material. Immediately after centrifugation, the clarified supernatants were transferred to fresh tubes and stored at-80°C until LC-MS analysis.

#### Tumor metabolites

Tumor metabolites were extracted using an organic solvent–based protocol. Briefly, 40–50 mg of mouse tumors (sample weights kept within a 1–1.5-fold range) were homogenized at full speed for one minute with a TissueLyser II (QIAGEN) containing frozen metal homogenization beads (7mm Stainless Steel beads, QIAGEN, #69990) and cold (-40°C) 70% (v/v) ethanol (LiChrosolv Ethanol gradient grade for LC, Supelco, #1.11727.1000). 14-fold volume of 75°C 70% (v/v) ethanol was added to the homogenized tumor samples. The samples were incubated for exactly one minute in a 75°C water bath, then vortexed quickly and transferred to an ice bath (<-20°C). The tubes were centrifuged 10 min at 4000 rpm at-4°C. The clarified supernatants were transferred to fresh tubes and stored at-80°C until LC-MS analysis.

#### LC-MS run parameters

Metabolome profiles of the sample extracts were acquired by General Metabolics using flow-injection mass spectrometry, as previously described (58). The instrumentation consisted of an Agilent 6550 iFunnel LC-MS Q-TOF mass spectrometer in tandem with an MPS3 autosampler (Gerstel) and an Agilent 1260 Infinity II quaternary pump. The running buffer was 60% isopropanol in water (v/v) buffered with 1 mM ammonium fluoride with Hexakis (1H, 1H, 3H-tetrafluoropropoxy)-phosphazene) (Agilent) and 3-amino-1-propanesulfonic acid (HOT) (Sigma Aldrich) as mass references. The isocratic flow rate was set to 0.150 mL/min. The instrument was run in 4GHz High Resolution, negative ionization mode. Mass spectra between 50 and 1,000 m/z were collected in profile mode. 5 uL of each sample were injected twice, consecutively, within 0.96 minutes to serve as technical replicates. The pooled study sample was injected periodically throughout the batch. Samples were acquired randomly within plates.

#### Data Processing & Annotation

Raw profile data were centroided, merged, and recalibrated using algorithms adapted from (58). Putative annotations were generated based on compounds contained in the Human Metabolome Database, KEGG, and ChEBI databases using both accurate mass per charge (tolerance 0.001 m/z) and isotopic correlation patterns.

#### Untargeted Metabolomics Analysis

Raw LC–MS data were first pre-processed (log10 transformed). The resulting feature table (a total of 1,642 metabolites; 208 samples from 75 mice) was normalized across all samples using the EigenMS method (R code available at https://sourceforge.net/projects/eigenms/files/R_EigenMS/) to reduce bias and technical variation (59).

Normalized data were stratified by tissue type (cecal content, serum, and tumor), each comprising seven groups (six antibiotic-treated groups and one control group). Principal component analysis (PCA) was performed separately for each tissue using R function stats::prcomp(X, center = TRUE, scale. = FALSE) to capture overall metabolic variation across samples. Group-level differences in metabolomic profiles were assessed by permutational multivariate analysis of variance (PERMANOVA) with 999 permutations (R vegan::adonis2) using Euclidean distances derived from PCA matrix. PCA plots were generated using the R package ggplot2 (cecal: Fig. 1D; serum: Fig. 1E; tumor: Fig. 1F).

EigenMS-normalized metabolite abundances were aligned by mouse ID to match cecal, serum, and tumor measurements from the same mice. For each metabolite, Spearman rank correlations were computed between cecal and serum, and between serum and tumor. Metabolites showing significant positive correlations in both comparisons after Benjamini–Hochberg correction (adjusted *P* < 0.05) were defined as putatively translocated from the gut to tumor tissue (n = 578). Tumor abundances of these 578 metabolites were then joined to log10-transformed tumor volume by mouse ID. Spearman correlations p-values were computed to quantify associations between metabolites and tumor volumes across mice.

### Targeted bile acid extraction and analysis

#### Serum bile acid extraction

A 50-μL aliquot of serum was added to a 96-well round-bottom polypropylene plate containing 100 μL of 1% (v/v) formic acid (Sigma-Aldrich, #27001-1L-R) and 20 μL of a custom-made stable isotope-labeled bile acid cocktail (Steraloids). The plate was vortexed for 30 s.

Subsequently, 350 μL of ice-cold acetonitrile (Honeywell, #HWLLC0154) was added, the plate was vortexed for an additional 30 s, and then allowed to stand for 5 min. The plate was centrifuged at 4500 rpm for 10 min.

#### Tumor bile acid extraction

Tumor tissue was placed into PowerBead Tubes (ceramic 2.8 mm beads, Qiagen, #13114-50) containing 1 mL of 70% (v/v) acetonitrile containing 0.2% (v/v) formic acid, and 20 μL of custom-made stable isotope-labeled bile acid cocktail (Steraloids). The samples were homogenized at 5.50 m s⁻¹ for 10 s using a Bead Ruptor 24 (Omni International). The homogenates were then centrifuged at 4500 rpm for 10 min.

#### Solid-phase extraction

Bile acids were isolated using solid-phase extraction with ISOLUTE PLD+ columns (50 mg/1 mL, Biotage, #918-0005-AG). The columns were sequentially conditioned with 2 mL of methanol (Fisher Scientific, #A456-4) followed by a 2 mL solution of 70% (v/v) acetonitrile containing 0.2% (v/v) formic acid. All solid-phase extraction steps were carried out on a positive-pressure manifold (Biotage) operated at 6 psi. The conditioned columns were placed into a 2-mL square 96-well polypropylene collection plate with 100-μL tapered reservoir (Analytical Sales, #968820). The supernatants were loaded onto the conditioned columns and eluted with two 500-μL aliquots of 70% (v/v) acetonitrile with 0.2% (v/v) formic acid. The eluates were collected using the positive-pressure manifold and evaporated to dryness under a gentle stream of nitrogen at 45°C. The dried residues were reconstituted in 500 μL of 50:50 (v/v) methanol/water with 0.2% (v/v) formic acid. The plate was vortex-mixed for 30 s and the samples were loaded for UPLC-MS/MS analysis.

Targeted bile acids were analyzed using ultra-high performance liquid chromatography coupled to tandem mass spectrometry (UPLC-MS/MS) (Shimadzu Nexera Series UPLC system coupled to a SCIEX Triple Quad 5500+ triple quadrupole mass spectrometer equipped with an electrospray ionization (ESI) source operated in negative ion mode). Chromatographic separation was performed on a Waters Acquity UPLC BEH C18 column (2.1 mm × 150 mm, 1.7 μm) using a gradient of Solvent A (25% Acetonitrile, 0.01% Formic Acid) and Solvent B (Acetonitrile 0.01% Formic Acid) at a flow rate of 0.35 mL/min. The column temperature was maintained at 45 °C. The gradient was: 0% to 20% B in 12 min; 20% to 74% B in 8 min; 74% to 100% B in 2.5 min, followed by re-equilibration at 0% B for 7.5 min. Quantification was performed using multiple reaction monitoring (MRM) with optimized Q1→Q3 transitions, collision energies, and dwell times as described in (60). Bile acid concentrations were quantified using calibration curves generated from authentic standards and normalized using stable isotope-labeled internal standards (**Table S5**). Peak integration and quantification were based on MRM peak areas. Data were processed to generate a metabolite concentration matrix for downstream analysis.

#### Metagenome Shotgun Sequencing and Analysis

Bacterial gDNA was extracted from fecal samples using the MagAttract Power Microbiome DNA/RNA KF kit (Qiagen, #27600-4-KF) on the Kingfisher Flex machine (ThermoFisher Scientific). DNA concentrations were quantified by a fluorescence-based assay (Quan-iT PicoGreen dsDNA, Invitrogen, #P11496). MSS was performed on an Illumina HiSeq 2000 (100-bp pair-end reads) at BGI Genomics.

Metagenomic shotgun sequencing (MSS) data from 69 samples were processed for taxonomic profiling using MetaPhlAn2 (61). Relative abundances of microbial taxa were obtained at the species level for downstream analyses.

Taxonomic relative abundances were summarized at the group level (six antibiotic-treated groups and one control group). Taxa were collapsed at multiple taxonomic levels (e.g., phylum, order, and family) for visualization. Stacked bar plots were generated to illustrate the relative abundance of major taxa across groups. Group-level abundances were calculated as the mean relative abundance across samples within each group.

Beta diversity was assessed using Bray-Curtis dissimilarity of species level bacterial relative abundances (R phyloseq::distance) (62). Principal coordinates analysis (PCoA) was performed using R function ape::pcoa to visualize differences in microbial community structure across groups, with group differences tested by PERMANOVA (R vegan::adonis2, 999 permutations) (63). The PCoA plot was generated using R ggplot2 (64).

#### Flow cytometry analysis

Tumor-bearing mice were euthanized (CO_2_ inhalation) at indicated time points, according to IACUC-approved protocols. Entire tumors (excess fat and lymph nodes were removed), spleens, and tumor draining lymph nodes (dLNs) were collected. Dissected tissues were kept in ice-cold sterile PBS until processing.

Single cells from dissected tumor tissue were isolated using the gentleMACS™ Octo Dissociator with Heaters (Miltenyi Biotec) and mouse Tumor Dissociation Kit (Miltenyi Biotec, #130-096-730), according to the manufacturer’s protocol. Briefly, the tumors were weighed and minced with a razor blade. The tumor pieces were mechanically and enzymatically dissociated using enzymes from the tumor dissociation kit and placed on the gentleMACS™ Octo Dissociator with Heaters (1x_m_impTumor_03 protocol) for 40 minutes. The dissociated tumor was then passed through a 70 µm MACS Smart Strainer (Miltenyi Biotec, #130-110-916) into a 15 mL conical tube and washed with 10 mL of RPMI medium (Gibco, #11875-119) with 10% FBS (Corning, #MT35015CV). The cell suspension was centrifuged at 400 x g, 4℃ for 10 min. The cell pellet was treated with RBC lysis buffer (Invitrogen, #00433357) to remove erythrocytes. Lymphocytes were isolated with a 40% Percoll solution (Cytiva, #17089101), centrifuged at 350 x g at RT for 20 min. The cell pellet was washed once with PBS prior to staining for flow cytometry.

Spleens and dLNs were mechanically dissociated by being mashed through a 70 μm sterile cell strainer (TWD Scientific LLC, #BX151070) in cold PBS. The cell suspensions were centrifuged at 400 x g, 4°C for 10 min. Splenocyte pellets were incubated with RBC lysis buffer (Invitrogen, #00433357) and washed in PBS. The single cell suspensions were kept in PBS, on ice, until flow cytometry staining.

Single-cell suspensions of cells were transferred to V-bottom 96 well plates (Corning, #3894) and centrifuged at 400 x g, 4℃ for 10 min. For surface staining, cells were incubated with Ghost 780 live/dead dye (Tonbo Biosciences) diluted in PBS (1:1000) at RT for 15 minutes. After washing with PBS, cells were incubated with α-CD16/CD32 blocking antibody (Biolegend), diluted 1:200 in FACS buffer (PBS, 3% heat-inactivated fetal bovine serum (Corning, #MT35016CV), 1 mM EDTA (Invitrogen, #AM9260G) for 10 min at 4℃ in the dark followed by surface staining with primary antibodies (1:200) for 30 min at room temperature in the dark. For surface staining, 10 μL of Horizon Brilliant Stain Buffer Plus (BD Biosciences, #566385) was added to each well to minimize staining artifacts. After staining, cells were washed with FACS buffer twice and suspended in FACS buffer prior to flow cytometry. Antibodies used for surface staining are listed in **Table S6**.

For intracellular staining, single cell suspensions were transferred to U-Bottom 96 well plates (Corning, #353077) and incubated in 200 μL RPMI supplemented with 10% FBS (Corning, #MT35015CV, 1x Cell Activation Cocktail (contains Phorbol 12-Myristate 13-Acetate, Ionomycin and Brefeldin-A; Biolegend, #423304) at 37°C with 5% CO2 for 4.5 hours. After incubation, the cells were washed with 1x PBS. The cells were incubated with 200 μL Ghost Dye Red 780 live/dead (Tonbo Biosciences) diluted in PBS (1:1000) at room temperature for 15 minutes. After washing with FACS buffer, cells were incubated with α-CD16/CD32 blocking antibody (Biolegend), diluted in FACS buffer 1:200, for 10 min at 4℃ in the dark followed by surface staining (as above).The cells were washed with FACS buffer twice and then fixed and permeabilized for 20 min at room temperature using the Cyto-fast Fix-Perm Buffer Set (Biolegend, #426803), according to the manufacturer’s instructions. The permeabilized cells were stained with antibodies targeting intracellular proteins (1:50) for 45 min at room temperature. Cells were washed with perm buffer twice and suspended in FACS buffer prior to flow cytometry. All flow cytometry experiments were performed on a Novocyte Advanteon (Agilent Technologies). Antibodies (clones and catalog numbers) used for flow cytometry are listed in **Table S6**.

#### Bulk cytokine/chemokine analysis from whole tumor lysates

Tumors were harvested from B16-F10 tumor-bearing mice. They were weighed and added to 200 µL of lysis buffer 2 (R&D Systems, #895347) supplemented with protease inhibitor tablets (Roche, #PIA32953) and phosphatase inhibitor tablets (Roche, #PIA32957). Tumor tissue lysates were prepared using a tissue homogenizer (TissueLyser II, Qiagen) and incubated on ice for 30 minutes. Samples were centrifuged at 15,000 RPM for 15 minutes at 4°C. The supernatant containing cytokines/chemokines was collected and stored in multiple aliquots at-80°C. Quantification of cytokines and chemokines was determined using the Mouse Luminex Discovery Assay (R&D Systems, #LXSAMSM) according to manufacturer’s instructions on a Bio-Plex 200 (Bio-Rad). Raw protein concentrations were normalized to tumor weight.

#### Single cell RNA sequencing of tumor-infiltrating immune cells

Single cell suspensions of tumors were prepared and stained with surface markers as described for flow cytometry. To account for individual variation, single cell suspensions from six mouse tumors of equivalent weight were pooled for each group (ICT + PBS and ICT + TDCA). Tumor-infiltrating immune cells (CD45^+^ cells) were sorted using a FACSymphony S6 cell sorter (BD Biosciences) into 1% FBS (Corning, #MT35015CV) in PBS. Sorted cellular suspensions of CD45^+^ cells were loaded onto a Chromium X (10X Genomics) at concentration of 1200 cells/µl to generate gel bead-in emulsions (GEM) (10X Genomics, Chromium Next GEM Chip G Single Cell Kit, #PN-1000120). Single cell RNA-seq libraries were prepared using the Chromium Single Cell 3’ v3.1 Gene Expression Kit (10x Genomics, #PN-1000269; Dual Index Kit TT Set A, 96 rxns #PN-1000215) according to manufacturer’s instruction. Each library was sequenced separately on a NextSeq 2000 (Illumina), using a P2-100 cycle flowcell, with at least 50,000 reads per cell. Library preparation and sequencing were performed by the UTSW McDermott Next Generation Sequencing Core.

Cell Ranger Software (v6.0) was used for sample demultiplexing, barcode processing, and single cell counting. Cell Ranger count was used to align samples to the reference mouse genome (UCSC mm10). The Seurat (v5.1.0) package in R (v4.3.1) was used for downstream analysis including clustering. Cells with mitochondrial content greater than 20% were removed from further analysis. In addition, cells with low Unique Molecular Identifier (UMI) (<6000) and gene number per cell (<100) were also excluded from further analysis. After filtering and integration(65), the dataset contained a total of 20,975 cells with a median UMI of 4,511 and median of 1,416 genes per cell. Uniform Manifold Approximation and Projection (UMAPs) were computed in Seurat using 30 dimensions. Cluster marker genes were identified using the FindAllMarkers function in Seurat, employing Wilcoxon rank-sum tests. Immune cell clusters were annotated based off canonical marker gene expression (See **Table S4**). Visualization of scatterplots were implemented using custom R scripts.

#### Statistical analysis

GraphPad Prism v.10 was used for statistical analyses. Data sets with normal distribution were analyzed with parametric tests, such as ordinary one-way ANOVA. For non-normal distributions, non-parametric tests, such as the Mann-Whitney U test or one-way ANOVA with Kruskal–Wallis test were applied. Survival was analyzed using the Mantel-Cox Log-rank test.

#### Language Editing Assistance

We used OpenAI’s ChatGPT (version 4.0 and 5.0) solely for language editing (e.g., grammar, clarity, and brevity) and for suggesting alternative phrasings during manuscript preparation. The tool did not contribute to study design, data generation, and analysis. It was not used to generate references. All suggestions from the ChatGPT were reviewed, revised, and approved by the authors. No text was inserted verbatim without subsequent human editing. The authors take full responsibility for the content of the manuscript.

