## Supplemental Figures for "A microbiota-derived bile acid overcomes antibiotic-induced hyporesponsiveness to immune checkpoint therapy by enhancing CD8^+^ T cell antitumor immunity"

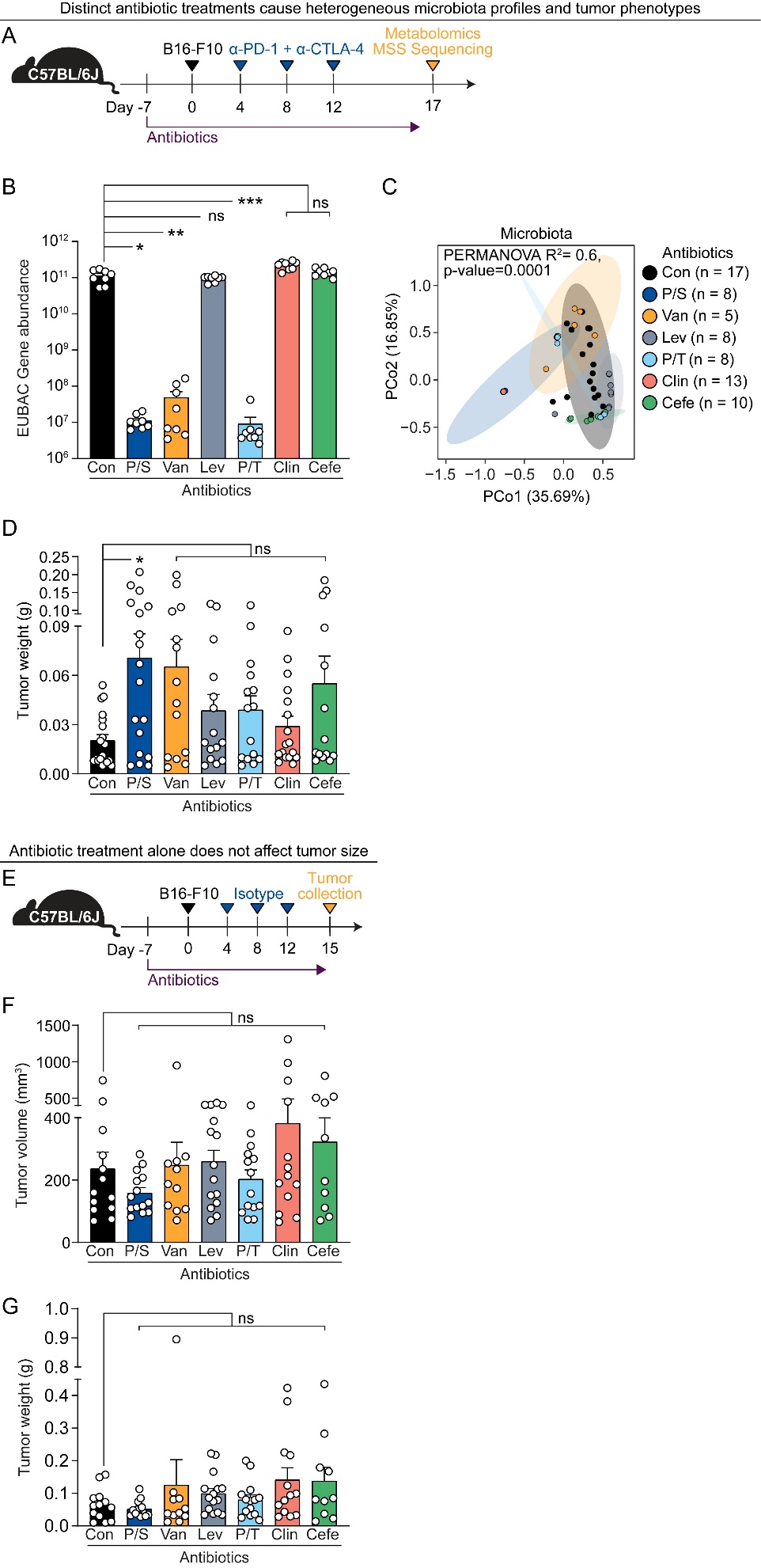

**Fig. S1. Antibiotic perturbation affects ICT efficacy and microbial communities.**

**(A)** Schematic overview of the B16-F10 heterotopic model, metabolomics and metagenomic shotgun sequencing (MSS). Mice (C57BL/6, female, 6–8 weeks old) were treated with different antibiotics throughout the experiment, starting one week before tumor inoculation: Con, no antibiotics control; P/S, Penicillin (0.95 g/L)/ streptomycin (2 g/L); Van, Vancomycin (0.5 g/L); Lev, Levofloxacin (0.5 g/L); P/T, Piperacillin/Tazobactam (12.5 mg/ml, I.P injection, once a day); Clin, Clindamycin (0.5 g/L); Cefe, Cefepime (12.5 mg/ml, I.P injection, twice a day). P/S, Van, Lev, Clin were administrated to mice via drinking water. 1 x 10^5^ B16-F10 cells were injected subcutaneously into the right inguinal flank. For ICT treatment, 200 µg of α-PD-1 and 200 µg α-CTLA4 (ICT) were injected intraperitoneally as shown. On day 17 post tumor inoculation, cecal content, serum and tumors were collected for metabolomics and stool was collected for MSS.

**(B)** Total bacterial DNA in stool from (A) was quantified by qPCR using universal 16S rRNA (EUBAC) primers. n = 5–17 mice per group.

**(C)** Beta-diversity analysis (Bray-Curtis distance with PERMANOVA) of microbiota in stool from (A). n = 5–17 mice per group.

**(D)** Tumor weights of ICT and antibiotic treated mice from (A), on day 17 post tumor inoculation. n = 14–19 mice per group.

**(E)** Schematic overview of the B16-F10 heterotopic model with isotype treatment. Mice (C57BL/6, female, 6–8 weeks old) were treated with different antibiotics throughout the experiment as in (A). 1 x 10^5^ B16-F10 cells were injected subcutaneously into the right inguinal flank. For ICT isotype treatment, 200 µg isotype control (rat IgG2a and mouse IgG2b) was injected intraperitoneally with the same timing as ICT treatment from (A).

**(F)** Tumor volumes of antibiotic treated mice from (E), on day 15 post tumor inoculation, with isotype treatment. n = 10–14 mice per group.

**(G)** Tumor weights of antibiotic treated mice from (E), on day 15 post tumor inoculation, with isotype treatment. n =10–14 mice per group.

ICT, immune checkpoint inhibitor therapy. Statistical analysis by one-way ANOVA, Kruskal–Wallis test with multiple comparisons for (B, D, F, G). Points represent individual mice. Bars denote mean ± SEM. All results are representative of ≥2 independent experiments. *p< 0.05; ****p** < 0.005; *****p** < 0.001; ns, not significant.

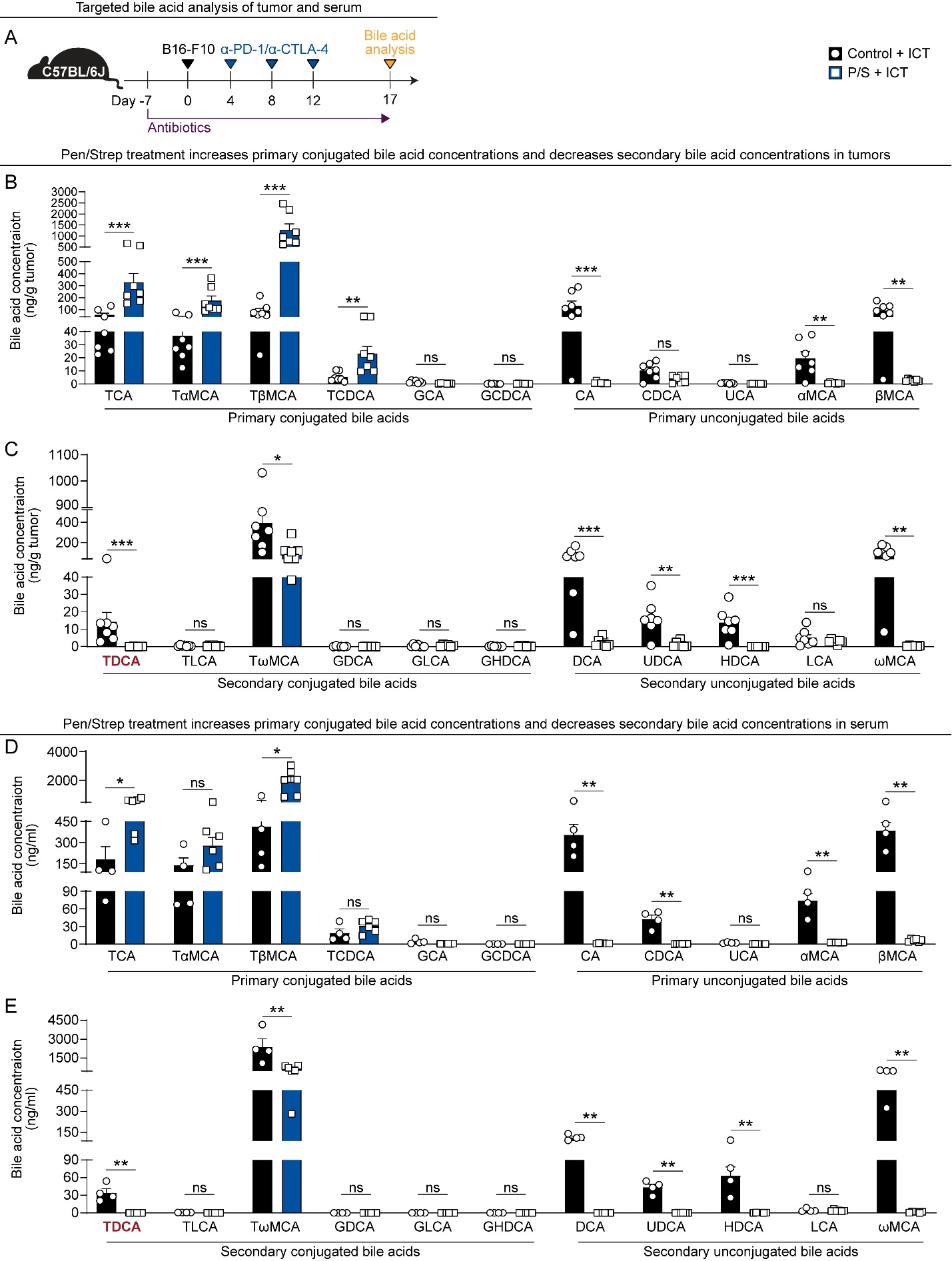

**Fig. S2. Pen/Strep treatment decreases secondary bile acids in the tumor and serum.**

**(A)** Schematic overview of targeted bile acid concentration determinations in tumor and serum. Mice were treated with antibiotics, B16-F10 cells, and ICT as in Figure 1A. Serum and tumors were collected on day 17 post tumor inoculation for targeted bile acid analysis by liquid chromatography-mass spectrometry (LC-MS). Con, no antibiotics control; P/S, Penicillin (0.95 g/L)/ Streptomycin (2 g/L).

**(B)** Concentrations of primary conjugated and primary unconjugated bile acids in tumors from (A).

**(C)** Concentrations of secondary conjugated and unconjugated bile acids in tumors from (A).

**(D)** Concentrations of primary conjugated and unconjugated bile acids in serum from (A).

**(E)** Concentrations of secondary conjugated and unconjugated bile acids in serum from (A).

TCA, taurocholic acid; TαMCA, tauro-α-muricholic acid; TβMCA, tauro-β-muricholic acid; TCDCA, taurochenodeoxycholic acid; GCA, [glycocholic acid](https://www.google.com/search?q=Glycocholic+acid&sca_esv=9506f15c4682863e&biw=2240&bih=1041&sxsrf=ANbL-n7fauK2W3B1Ije94Ass6xQ6nr0YUw%3A1775158391749&ei=d8TOabS6LZynmtkP3cGC4Qg&ved=2ahUKEwjVs9Cw9M-TAxV2nGoFHTXvMC4QgK4QegQIARAC&uact=5&oq=bile+acid+GCA+full+name&gs_lp=Egxnd3Mtd2l6LXNlcnAiF2JpbGUgYWNpZCBHQ0EgZnVsbCBuYW1lMgUQABjvBTIFEAAY7wUyBRAAGO8FSJ00UMYIWKEycAV4AZABAJgBWqAB-gaqAQIxNLgBA8gBAPgBAZgCE6ACzAfCAgoQABhHGNYEGLADwgIEEAAYHsICCxAAGIAEGIoFGIYDwgIHEAAYgAQYDcICBhAAGB4YDcICCBAAGAUYHhgNwgIIEAAYCBgeGA3CAggQABiABBiiBMICCBAAGIkFGKIEmAMAiAYBkAYIkgcCMTmgB_MosgcCMTS4B7sHwgcGMC4xMi43yAczgAgB&sclient=gws-wiz-serp&mstk=AUtExfDG9fjP3ED8QKgYMoJnpWDu9mGyezAwaKkfsKkr7vJ1sjbt12PG6cZ2YXK2hMmUmU79yZzC2Gd63wr69md2jCyp9w8Y5626mARXmafHVrRaSxnabsc_fodSZ8y5Q7tc9BVtBlCEkVrCjUb2zTNDe1swxqwVw0Dnegz3IEXqghwptYceXJqHDobxQDiyr3wkH20GE_n81xY1Ewr20PLUzAKFMned0qwuNFNxFjFZU4_zYuOXIbTU5feE3JbPelv1yxAjJQoKbCaA--y0g2oN8qFP&csui=3); GCDCA, [glycochenodeoxycholic acid](https://www.google.com/search?q=Glycochenodeoxycholic+acid&sca_esv=9506f15c4682863e&biw=2240&bih=1041&sxsrf=ANbL-n725Efr_jS8GRtgGygUAqFVs0Je-w%3A1775158403580&ei=g8TOafuOI_qmmtkPqJWy6QM&ved=2ahUKEwiO0uTA9M-TAxXVnWoFHVUAK-IQgK4QegQIARAC&uact=5&oq=bile+acid+GCDCA+full+name&gs_lp=Egxnd3Mtd2l6LXNlcnAiGWJpbGUgYWNpZCBHQ0RDQSBmdWxsIG5hbWUyCBAhGKABGMMEMggQIRigARjDBEjKClCCAliCAnABeAGQAQCYAUigAUiqAQExuAEDyAEA-AEC-AEBmAICoAJgwgIKEAAYRxjWBBiwA5gDAIgGAZAGCJIHATKgB6ICsgcBMbgHTsIHBzAuMS4wLjHIBwyACAE&sclient=gws-wiz-serp&mstk=AUtExfB7yQrDCRx6ASIxKRl59M1xqc6TlsliXhpLVK8rGqpMrNKOInM1scmMRkFmP6AdYLsb_uGk33aT1s33ZTFHMInVePLkKpk1ooxkPX-KrJGvmIo71Q6_XOwcMmAzLpCrFNVvfruM3EoBFSn07VdEUUeb_XBpQDpOdL6aeaotj_sVeASFhG9BF_Q67aBtJiqqJO4sHuuzo9GtaSTSyAzSzAJqbIXcgnrDoGmG5l21hQfLX0psIw28NfAVRfu1ZL8DAYVcinQq_wLy8h-Eh5JrhEjF&csui=3); CA, [cholic acid](https://www.google.com/search?q=Cholic+acid&sca_esv=9506f15c4682863e&biw=2240&bih=1041&sxsrf=ANbL-n6Ua2YGklMYuAwQDw4909aTeSLiPw%3A1775158437427&ei=pcTOaZnXGdmkqtsPk-_CYA&ved=2ahUKEwiKtMHM9M-TAxXMkGoFHWxMAQQQgK4QegQIARAE&uact=5&oq=bile+acid+CA+full+name&gs_lp=Egxnd3Mtd2l6LXNlcnAiFmJpbGUgYWNpZCBDQSBmdWxsIG5hbWUyCBAhGKABGMMESPMRUMQHWI0PcAF4AZABAJgBU6ABuQGqAQEzuAEDyAEA-AEBmAIEoALTAcICChAAGEcY1gQYsAPCAgUQABjvBcICBBAjGCeYAwCIBgGQBgiSBwE0oAfeBbIHATO4B8sBwgcFMC4xLjPIBw-ACAE&sclient=gws-wiz-serp&mstk=AUtExfAYsAEIZuBniIBlFNmneaCddwuxyyZwL9C2dDgu7M8SLHCojfi7SvpfaH2MQBtDjuegcMgZ-rupiGNz3yMxdNPWHJ9D_kMK6HEq_WKu-AvpXXlzXabot2XrLJUvP9jex8xtgbUQtlgGNHnvDmsqV_ggSVfsj2b-m-OIOTDZeyHwBYlm_2UoRJNvx4kc1541tQqDCGF68rJVOuF0UZOv6-hwyL6yiqqggsO7mpA0Xcx1CxwxTgqETLopLr0rswAtqODx133_v0D7AUQrw02sJ4sB&csui=3); CDCA, chenodeoxycholic acid; UCA, [ursocholic acid](https://www.google.com/search?q=Ursocholic+acid&sca_esv=9506f15c4682863e&biw=2240&bih=1041&sxsrf=ANbL-n6zUGi_bDkwH0t0tYqa98fhW5WFQw%3A1775158481384&ei=0cTOaaiNF-2gqtsPrKjBwQk&ved=2ahUKEwiQr4re9M-TAxV5l2oFHWZBJl4QgK4QegQIARAB&uact=5&oq=bile+acid+UCA+full+name&gs_lp=Egxnd3Mtd2l6LXNlcnAiF2JpbGUgYWNpZCBVQ0EgZnVsbCBuYW1lMggQIRigARjDBEjEDVDnAVilB3ABeAGQAQCYAUagAX2qAQEyuAEDyAEA-AEC-AEBmAIDoAKSAcICChAAGEcY1gQYsAPCAgoQIRgKGKABGMMEmAMAiAYBkAYIkgcBM6AHuASyBwEyuAeHAcIHBzAuMi4wLjHIBwyACAE&sclient=gws-wiz-serp&mstk=AUtExfCzCvlks7pWgZ_ICJDqP_P4ld63BYVUFVs4EsE561AT63Tk6qEJQH7-Fp-GRaMqRMevQUJ15UfHBDQzDnaZjljiXnLrLln-TByH0vnujLH3YBF02Nqqm3ywB7rhCqy-UIWZAAaSdijSlDjVp5sL9D32NNRARqOL_7Hq_vziNCRBBZDLgqcoHhoC5mJwfCWQXrlH8lfn_2p_DWv-f8GGg6-wKEPhvXnlB2pqWST9Owma9dqueu_1kmNVfFN1tTZ5y64vDCpTwvqbTGyW2Q7KxQWM&csui=3); αMCA, α-muricholic acid; βMCA, β-muricholic acid; TDCA, taurodeoxycholic acid; TLCA, taurolithocholic acid; TωMCA, [tauro-ω-muricholic acid](https://www.google.com/search?q=Tauro-%CF%89-muricholic+acid&sca_esv=9506f15c4682863e&biw=2240&bih=1041&sxsrf=ANbL-n7njNLN87hl5EGB_Xt0TB5xcOxsHA%3A1775158542046&ei=DsXOaZe3AvitqtsPpp6CkQc&ved=2ahUKEwjd95mD9c-TAxVVkWoFHZkXL-AQgK4QegQIARAC&uact=5&oq=bile+acid+T%CF%89MCA%2C+full+name&gs_lp=Egxnd3Mtd2l6LXNlcnAiG2JpbGUgYWNpZCBUz4lNQ0EsIGZ1bGwgbmFtZTIIEAAYgAQYogQyBRAAGO8FMgUQABjvBTIFEAAY7wVIzwtQAFgAcAB4AZABAJgBSqABSqoBATG4AQPIAQD4AQL4AQGYAgGgAlGYAwCSBwExoAeJArIHATG4B1HCBwMyLTHIBwWACAE&sclient=gws-wiz-serp&mstk=AUtExfDO6oHPQFIkmXYVsVJOg0A_Owp2TQtStJ4VKb04SIfGo_JLXt88EALeBDniR0sfGSWT7YrZxgid0iG8Cynerzh7tX7ahGAxWdJFkzh7ExBw12vyZpP00-MGXrvEZjNFkUMNzIzieQtYsYt8Pii-yQGPlfOscghfQzzCtgiNTAmUdxzaB7qiPKhYFfsJPJdYDnQ-1lHqE14-wqwkTWNoSnmgq7_fAZFYB1aGN324hvkerYbdehV29NTFWDh_iss1M0k2x3A6l7ZTu7kZdEwMdO-k&csui=3); GDCA,  [glycodeoxycholic acid](https://www.google.com/search?q=Glycodeoxycholic+acid&sca_esv=9506f15c4682863e&biw=2240&bih=1041&sxsrf=ANbL-n54AwqfXtFa1fxla1MCC5FErzHc8g%3A1775158576639&ei=MMXOadDUJoSyqtsPt6bzkQw&ved=2ahUKEwi2ovCN9c-TAxUy5ckDHSuPNWYQgK4QegQIARAC&uact=5&oq=bile+acid+GDCA+full+name&gs_lp=Egxnd3Mtd2l6LXNlcnAiGGJpbGUgYWNpZCBHRENBIGZ1bGwgbmFtZTIIECEYoAEYwwRIxQtQnQJYnQJwAXgAkAEAmAFIoAFIqgEBMbgBA8gBAPgBAvgBAZgCAaACTpgDAIgGAZIHATGgB6wBsgcBMbgHTsIHAzAuMcgHAoAIAQ&sclient=gws-wiz-serp&mstk=AUtExfAxHqzbWPWhix5aWA7v3RUxFl0narEW2tQAyD7AWuANxVeHMZkyBV15K-HAwUCbKM-O2YQWXlVvsMgwu4iqGHKRlVDDidNSfkGisWKRIr-zTNct3rx_EUV4o3GX_eGm4zdBjDm7u9jvBZseajcysPERvx2KYT7pbhLlJB5rObNARdB9T5G6jIQprk59gAN42VaPi3K7kandYPeJ93bmymomdUCn62vw4FgBSWV1-Yd-sOAH07QHjcntiEJ5WJ1sYiy8qiS1PJdl6gf8Pvc5k0cj&csui=3); GLCA, [glycolithocholic acid](https://www.google.com/search?q=Glycolithocholic+acid&sca_esv=9506f15c4682863e&biw=2240&bih=1041&sxsrf=ANbL-n7-avrzgIZ81j5L5hYY9XieQ3zfIg%3A1775158599127&ei=R8XOaaa_B-21qtsP6r-OkAY&ved=2ahUKEwiL4rKa9c-TAxWgl2oFHRZXM6IQgK4QegQIARAC&uact=5&oq=bile+acid+GLCA+full+name&gs_lp=Egxnd3Mtd2l6LXNlcnAiGGJpbGUgYWNpZCBHTENBIGZ1bGwgbmFtZTIFEAAY7wUyBRAAGO8FMgUQABjvBTIFEAAY7wVI_glQugFYugFwAXgBkAEAmAFLoAFLqgEBMbgBA8gBAPgBAvgBAZgCAqACXcICChAAGEcY1gQYsAOYAwCIBgGQBgiSBwEyoAfOAbIHATG4B03CBwcwLjEuMC4xyAcLgAgB&sclient=gws-wiz-serp&mstk=AUtExfB_5D59UYto42RmGI0GOTSVbN0c6vn2bg8S2yL8Y9IXoStlvNaQy02GcenPfkevW5JmS5Pphp6b81FkJoMfcAoegTlU0c6nP54LV9dy9lhIeHMQNFCdX2tBL6Bp7EiQiquuHr9pedKiFGxrFT8taHNcNQFKuBxyyczDIHiACZCZuEBbLItw_ZW9fvS05qSN2GcxwB5XQ4hdIA2xwkHwSKtYQuC50mrTFI25SY2MkSgjCIGqQwBAm7vAXTpok7Vh58js-o6ixEv5bJ0-k7bbbvP_&csui=3); GHDCA, [glycohyodeoxycholic acid](https://www.google.com/search?q=Glycohyodeoxycholic+Acid&sca_esv=9506f15c4682863e&biw=2240&bih=1041&sxsrf=ANbL-n6fQA-tTK8TSzhFnrni9y-I__tafA%3A1775158625401&ei=YcXOaZmSGKelqtsPnujB0AM&ved=2ahUKEwiSifOi9c-TAxXAkmoFHcRgG4sQgK4QegQIARAC&uact=5&oq=bile+acid+GHDCA+full+name&gs_lp=Egxnd3Mtd2l6LXNlcnAiGWJpbGUgYWNpZCBHSERDQSBmdWxsIG5hbWUyBxAjGLACGCcyCBAAGIAEGKIEMgUQABjvBTIFEAAY7wUyBRAAGO8FMgUQABjvBUjnC1DkBFjkBHABeAGQAQCYAUegAUeqAQExuAEDyAEA-AEC-AEBmAICoAJXwgIKEAAYRxjWBBiwA5gDAIgGAZAGBJIHATKgB_QEsgcBMbgHS8IHAzItMsgHCoAIAQ&sclient=gws-wiz-serp&mstk=AUtExfB9Q9XGGD3PVhhk89LtsbB9y2fUGUB1xSAesuyF0JRW5yQqR_lMbXsMoaS2JeqqtvCiJetF7jSZJ_6QBzmz1Ta990OibiCF1GJMBSQEgZHXs3QWVeiMvmnqNdmCwvmwimY7OxysJMtoXb589Krn9SWVUBGV7iLAS9cqxuNcs7wEIpSECE_LPlaYFKW2tXvgiWrm5U59B8cuZtZdKEWwr3X4x0fe_tlb_ogU_2TlwLitml337AVV67CwSMvCPQ1__euI9Tzygbkzron7Nr-cU6nt&csui=3); DCA, deoxycholic acid; UDCA, [ursodeoxycholic acid](https://www.google.com/search?q=Ursodeoxycholic+acid&sca_esv=9506f15c4682863e&biw=2240&bih=1041&sxsrf=ANbL-n55RvuLrq63O_Nd2i5VY1NaQ9Mm7g%3A1775158704457&ei=sMXOac-_G5KqqtsPyPbcgAs&ved=2ahUKEwiB1rXP9c-TAxXtlCYFHX5KODwQgK4QegQIARAC&uact=5&oq=bile+acid+UDCA+full+name&gs_lp=Egxnd3Mtd2l6LXNlcnAiGGJpbGUgYWNpZCBVRENBIGZ1bGwgbmFtZTIIECEYoAEYwwRIvQhQAFgAcAB4AZABAJgBSaABSaoBATG4AQPIAQD4AQL4AQGYAgGgAlKYAwCSBwExoAesAbIHATG4B1LCBwMwLjHIBwKACAE&sclient=gws-wiz-serp&mstk=AUtExfClX8CvA3qet9vCucLRxGKdO9K1gCcjXNUu7HrB0tQ9DHTklaS3FDZhUDwe1buqMHnQpIpF57XzPJ__6mzSCxalE8qfyH4SiOsBWhnC4CYkbRBezEzj6x2aIwcT0LhWvoLud78qfZmdLaqewGcHeFAd_xj3jrCMJJjWkp57OSQc-Dg_lS-DWoCEm5oZhVYmNKuhi0PpPS4h_Ibya9MhxOLBIT1zSogTNL0ShGDWwY7ucAegRdXgzKCo4wRevt4q_9FsLpPTOfo7PP-EQ6J7JvI_&csui=3); HDCA, [hyodeoxycholic acid](https://www.google.com/search?q=Hyodeoxycholic+acid&sca_esv=9506f15c4682863e&biw=2240&bih=1041&sxsrf=ANbL-n4K2aezqEjJlveK0q5JINQsO-21jA%3A1775158736620&ei=0MXOaenAJbK7qtsPg6bRyAw&ved=2ahUKEwi9kYba9c-TAxV-nCYFHa0ANaAQgK4QegQIARAC&uact=5&oq=bile+acid+HDCA%2C++full+name&gs_lp=Egxnd3Mtd2l6LXNlcnAiGmJpbGUgYWNpZCBIRENBLCAgZnVsbCBuYW1lMgUQABjvBTIFEAAY7wUyBRAAGO8FSIQLUABYAHAAeAGQAQCYAUigAUiqAQExuAEDyAEA-AEC-AEBmAIBoAJTmAMAkgcBMaAHkQGyBwExuAdTwgcDMy0xyAcJgAgB&sclient=gws-wiz-serp&mstk=AUtExfAbjWzPtGGK-s4lgVA2hSXsuHWmupVdnhN7BZ3cLczPHxFQs-b5hWT0eEIrgC5IXxCWhit1k-UJsjgczTngAdSok3Amn_upYOIoKw9ANNXN4LhHolZPYkf9lBkABxohPILbssVz_um8IKcfuqRW2m2W1HGGQflAmMtzcZ6ObstgJzLHRTDhMPy_OL8eI6q22WFXGExqZAGE3bcqCQe2EOUecW2spgLIpJkRzaIPY2dvprf_HJB5ZOGif9NVh0QUWdqtH9m-ngq9O-bD2zpFuIn1&csui=3); LCA, [lithocholic acid](https://www.google.com/search?q=Lithocholic+acid&sca_esv=9506f15c4682863e&biw=2240&bih=1041&sxsrf=ANbL-n4spUIDh9j1ZG9cjgPRBGJCTkmRqw%3A1775158643207&ei=c8XOaaqpDImtqtsPzqnMiQ0&ved=2ahUKEwioj4zA9c-TAxXEmSYFHW3dEIQQgK4QegQIARAC&uact=5&oq=bile+acid+LCA+full+name&gs_lp=Egxnd3Mtd2l6LXNlcnAiF2JpbGUgYWNpZCBMQ0EgZnVsbCBuYW1lMggQIRigARjDBEjPC1DMAVjMAXABeAGQAQCYAYABoAGAAaoBAzAuMbgBA8gBAPgBAvgBAZgCAqACmAHCAgoQABhHGNYEGLADmAMAiAYBkAYIkgcDMS4xoAepAbIHAzAuMbgHhAHCBwcwLjEuMC4xyAcNgAgB&sclient=gws-wiz-serp&mstk=AUtExfDxYtF_OAOdPUl29KTE4bhACygm7C1Eq_uTvqjSBetKXlwuP1k58JjIl40D7B6lJ0wlsktDXx-Z9JRDSZO_LjuAkX-9zqxJDnUS8dIiLIXIcAnWH5vqOe1PbaRYSYwlgDUts7pHBHGaqPKphkQBNhrzqD-5e7lpzWUkmgC5PipXwPZzDP925IDa-niKaeQ-V0BIKrpjvRK188tvAxfI4uMweZz1dDRp0KgNLg-PXM03tHD-0SA_DAz9wt_VlhcLI9mx8bVKfuLefS3YcQUcpB3c&csui=3); ω-MCA, ω-muricholic acid. Statistical analysis by Mann–Whitney test for (B–E). Points represent individual mice. Bars denote mean ± SEM. All results are representative of ≥2 independent experiments. *p< 0.05; ****p** < 0.005; *****p** < 0.001; ns, not significant.

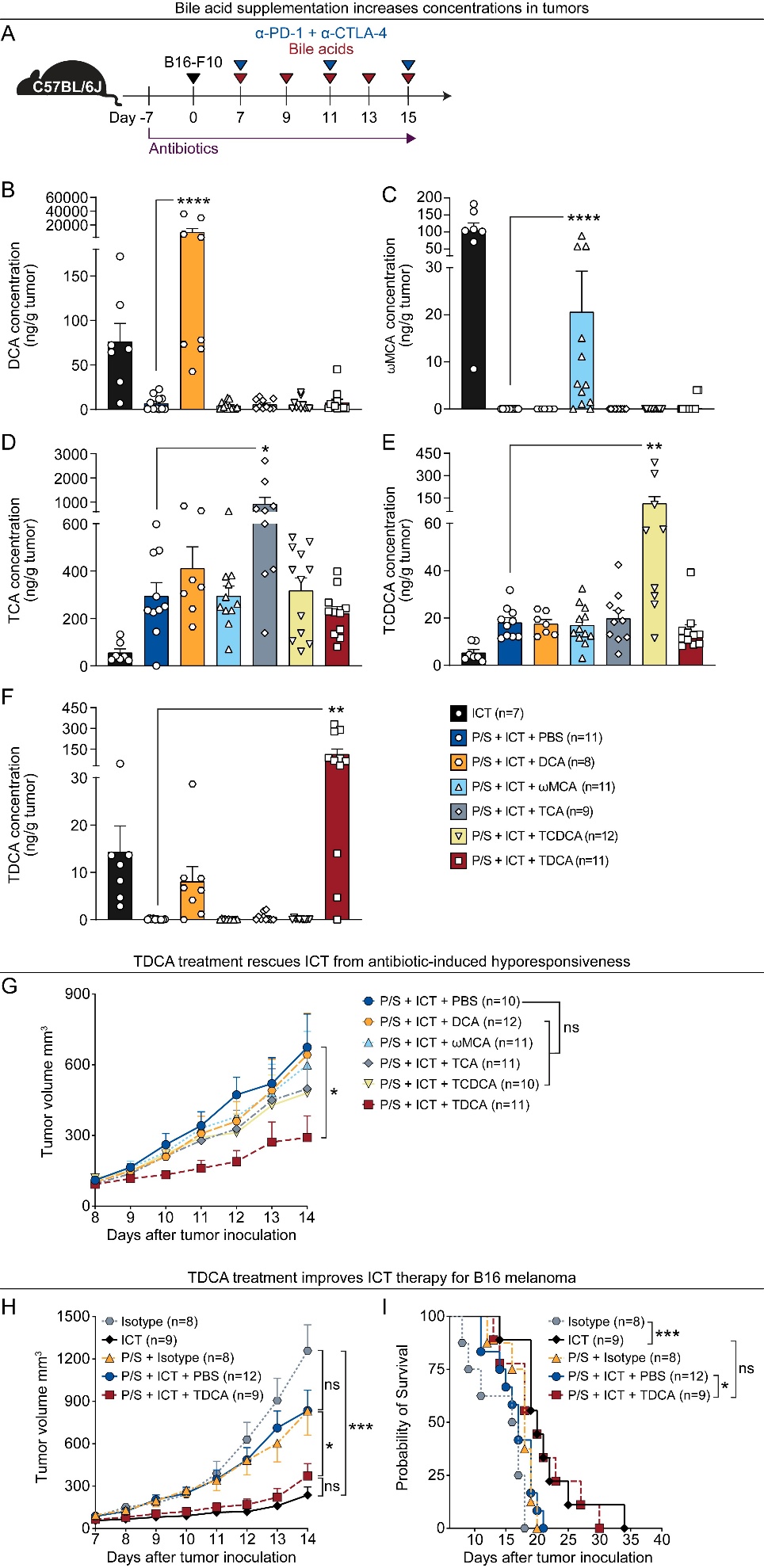

**Figure S3. Only TDCA supplementation rescues antibiotic-induced ICT hyporesponsiveness.**

**(A)** Schematic overview of the B16-F10 heterotopic model with TDCA treatment. Mice (C57BL/6, female, 6–8 weeks old) received penicillin/ streptomycin (0.95 g/L; 2 g/L) in drinking water throughout the experiments. 1 x 10^5^ B16-F10 cells were injected subcutaneously into the right inguinal flank. ICT or isotype control was injected intraperitoneally, beginning when tumor volumes reached 80 ± 20 mm³. TDCA (50 μl of 4 mM), DCA (50 μl of 4 mM), TCA (50 μl of 16 mM), TCDCA (50 μl of 8 mM), ω-MCA (50 μl of 2 mM) were administrated every two days by intratumoral injection.

**(B-F)** Quantification of bile acids in tumors from (A) by liquid chromatography-mass spectrometry (LC-MS) on day 16 post tumor inoculation. DCA **(B)**, ω-MCA **(C)**, TCA **(D)**, TCDCA **(E)** and TDCA **(F)** n =7–12 mice per group.

**(G)** Tumor volumes following ICT and treatment of different bile acids (TDCA, DCA, TCA, TCDCA, ω-MCA) as described in (A); n =10–12 mice per group.

**(H)** Tumor volumes following ICT and antibiotic treatment with or without TDCA treatment (50 μl of 4 mM) as described in (A); n =8–12 mice per group.

**(I)** Survival of tumor-bearing mice ICT and antibiotic treatment with or without TDCA treatment as described in (A); n = 8–12 mice per group.

DCA, deoxycholic acid; ω-MCA, ω-muricholic acid; TCA, taurocholic acid; TCDCA, taurochenodeoxycholic acid; TDCA, taurodeoxycholic acid. Statistical analysis by one-way ANOVA, Kruskal–Wallis test (B-H) or Log-rank test for (I). Points represent individual mice. Bars denote mean ± SEM. All results are representative of ≥2 independent experiments. *p< 0.05; ****p** < 0.005; *****p** < 0.001; ns, not significant.

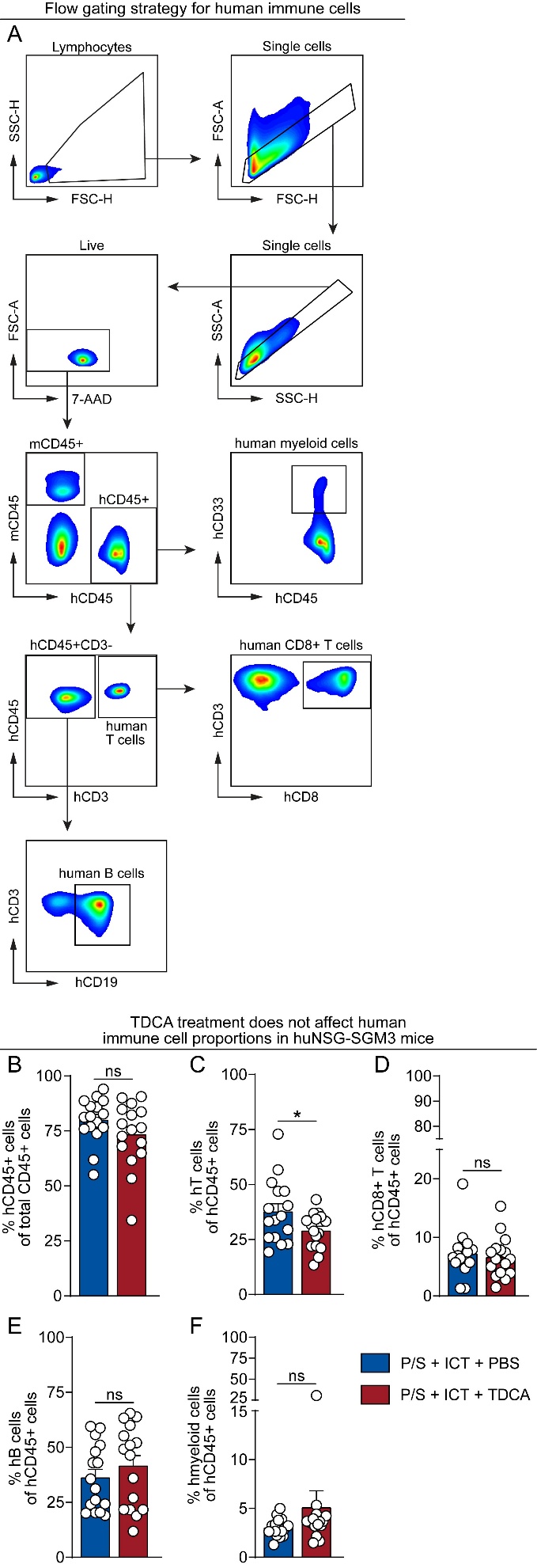

**Figure S4. huNSG-SGM3 mice have successful engraftment of hCD34^+^ cells.**

**(A-F)** NSG-SGM3 mice (4–8 weeks old, female) were irradiated with 240-250 cGy 24 hours before tail vein transfer of 1 x 10^5^ human hematopoietic CD34^+^ cord blood cells. After 10 to 12 weeks, human immune system engraftment was evaluated by flow cytometry of peripheral blood.

**(A)** Flow cytometry gating strategy for human immune cells in huNSG-SGM3 mice.

**(B)** Percentage of hCD45^+^ cells out of total CD45^+^ cells in peripheral blood.

**(C-F)** Percentage of human immune cells out of hCD45^+^ cells in peripheral blood: T cells **(C)**, CD8⁺ T cells **(D)**, B cells **(E)**, myeloid cells **(F)**. n = 16 mice per group.

huNSG-SGM3, humanized NSG-SGM3 mice; hCD45^+^, humanized CD45^+^. Statistical analysis by Mann–Whitney test (C-K). Points represent individual mice. Bars denote mean ± SEM. All results are representative of ≥2 independent experiments.  *p< 0.05; ns, not significant.

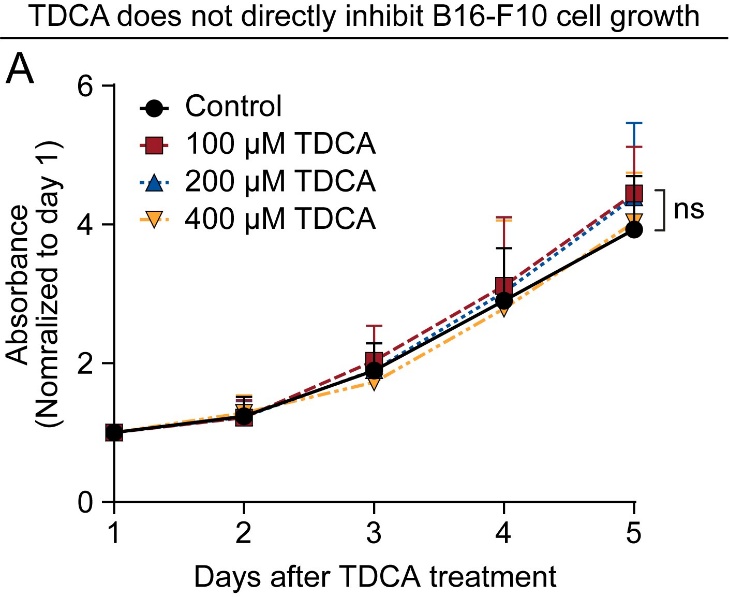

**Figure S5. TDCA does not directly inhibit B16-F10 cell growth or cause cell death.**

**(A)** B16-F10 cells were treated with increasing concentrations of TDCA (0 μM, 100 μM, 200 μM, 400 μM), and cell viability was assessed using an MTS assay. Absorbance values were measured and normalized to day 1.

TDCA, taurodeoxycholic acid. Statistical analysis by one-way ANOVA, Kruskal–Wallis test. ns, not significant.

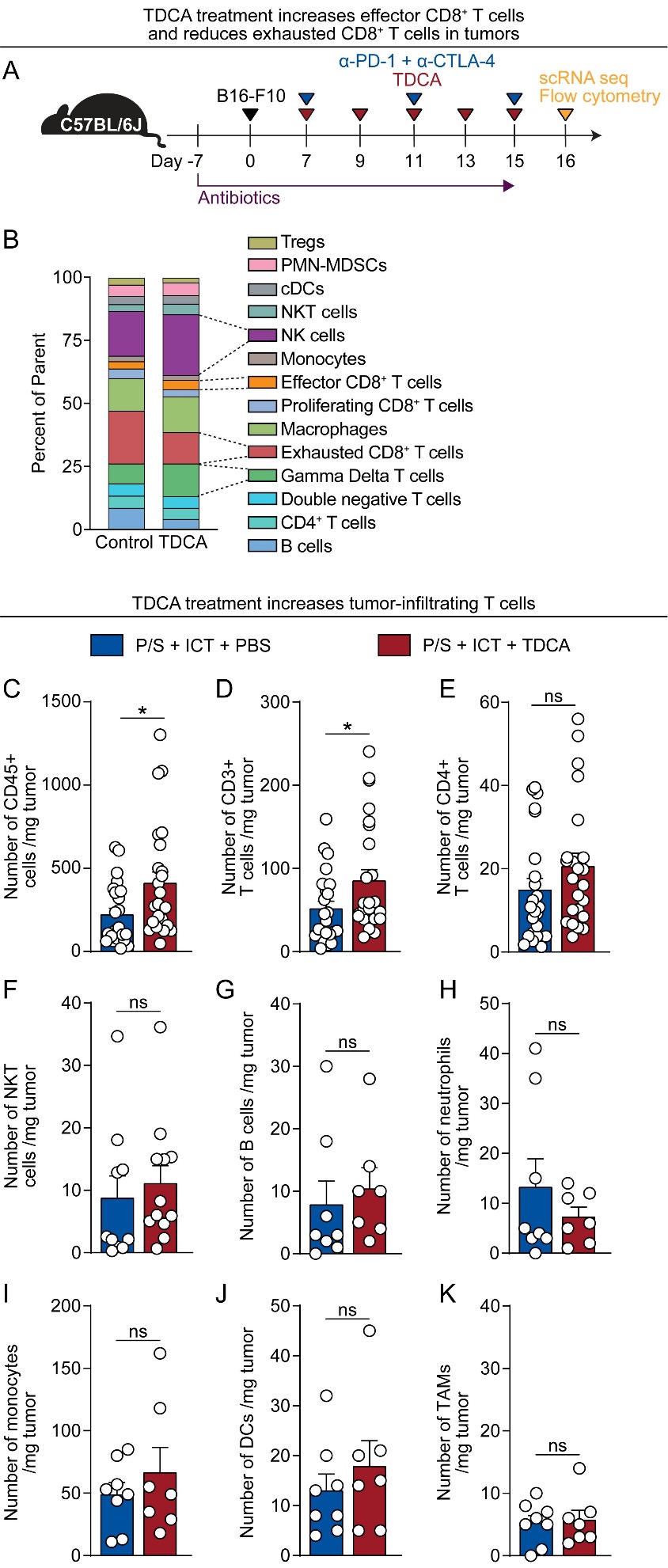

**Figure S6. TDCA treatment increases effector CD8^+^ T cells and NK cells in tumors.**

**(A)** Schema of tumor immune profiling experiments. Mice received penicillin/ streptomycin (0.95 g/L; 2 g/L) in drinking water throughout the experiment. 1 x 10^5^ B16-F10 cells were injected subcutaneously into the right inguinal flank. ICT was injected intraperitoneally, beginning when tumor volumes reached 80 ± 20 mm³. TDCA (50 μl of 4 mM) was intratumorally injected every two days. Tumors were collected on day 16 post inoculation for immune profiling.

**(B)** Proportions of cell clusters identified by single-cell RNA sequencing.

**(C-K)** Quantification of tumor-infiltrating immune cells by flow cytometry. CD45^+^ cells **(C)**, CD3⁺ T cells **(D)**, CD4⁺ T cells **(E)**, natural killer T cells **(F)**, B cells **(G)**, neutrophils **(H)**, monocytes **(I)**, dendritic cells **(J)**, and tumor-associated macrophages (**K**). n = 7–25 mice per group.

scRNA seq, single-cell RNA sequencing; TDCA, taurodeoxycholic acid; NK cells, natural killer cells; NKT, natural killer T cells; DCs, dendritic cells; TAMs, tumor-associated macrophages. Statistical analysis by Mann–Whitney test (C-K). Points represent individual mice. Bars denote mean ± SEM. All results are representative of ≥2 independent experiments. *p< 0.05; ns, not significant.

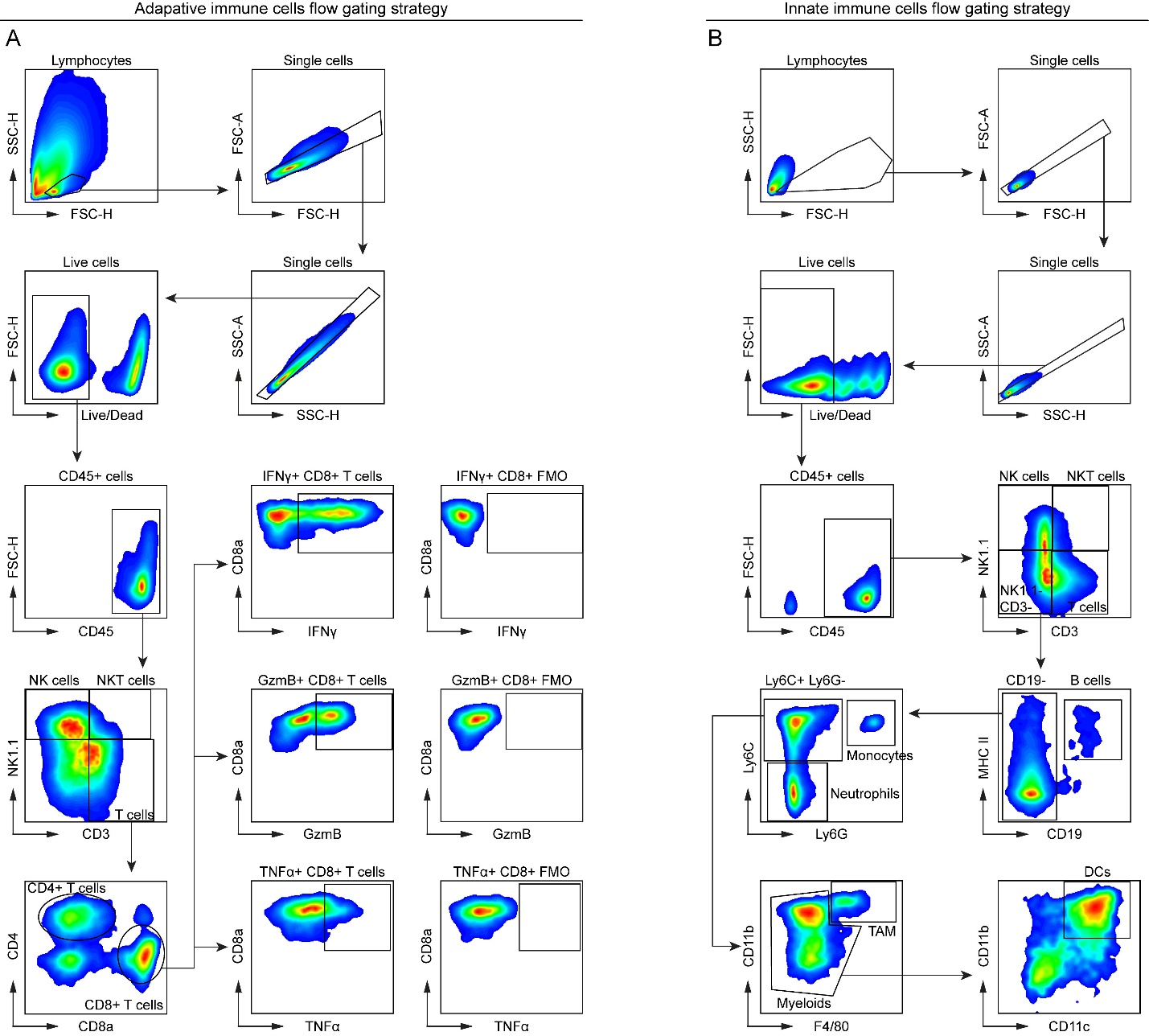

**Figure S7. Flow cytometry gating strategy for tumor-infiltrating immune cells and immune cells in the draining lymph node (dLN).**

**(A)** Flow cytometry gating strategy for adaptive immune cells.

**(B)** Flow cytometry gating strategy for innate immune cells.

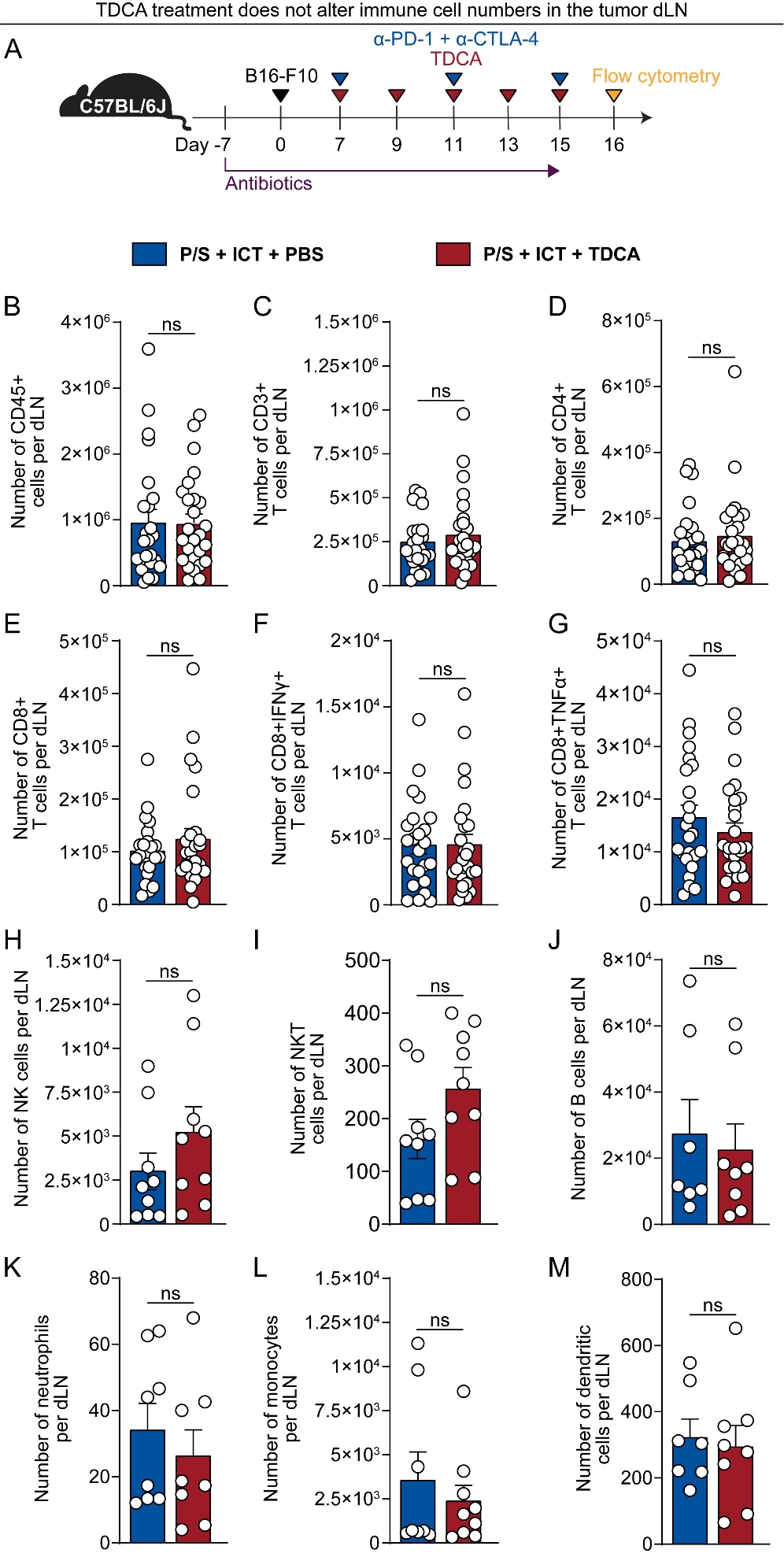

**Figure S8. Immune cell populations in the tumor draining lymph node are not affected by TDCA treatment.**

**(A)** Schematic of immune profiling of tumor draining lymph nodes by flow cytometry. Mice received penicillin/ streptomycin (0.95 g/L; 2 g/L) in drinking water throughout the experiment. 1 x 10^5^ B16-F10 cells were injected subcutaneously into the right inguinal flank. ICT or isotype control was injected intraperitoneally, beginning when tumor volumes reached 80 ± 20 mm³. TDCA (50 μl of 4 mM) was intratumorally injected every two days. Tumors were collected on day 16 post inoculation for immune profiling.

**(B-M)** Quantification of immune cell populations in dLN by flow cytometry. n = 7–25 mice per group.

**(B)** CD45^+^ cells

**(C)** CD3⁺ T cells

**(D)** CD4⁺ T cells

**(E)** CD8⁺ T cells

**(F)** IFN-γ⁺ CD8⁺ T cells

**(G)** TNF-α⁺ CD8⁺ T cells

**(H)** NK cells

**(I)** NKT cells

**(J)** B cells

**(K)** Neutrophils

**(L)** Monocytes

**(M)** Dendritic cells

dLN, draining lymph node; TDCA, taurodeoxycholic acid; NK cells, natural killer cells; NKT, natural killer T cells. Statistical analysis by Mann–Whitney test for (B-M). Points represent individual mice. Bars denote mean ± SEM. All results are representative of ≥2 independent experiments.  ns, not significant.

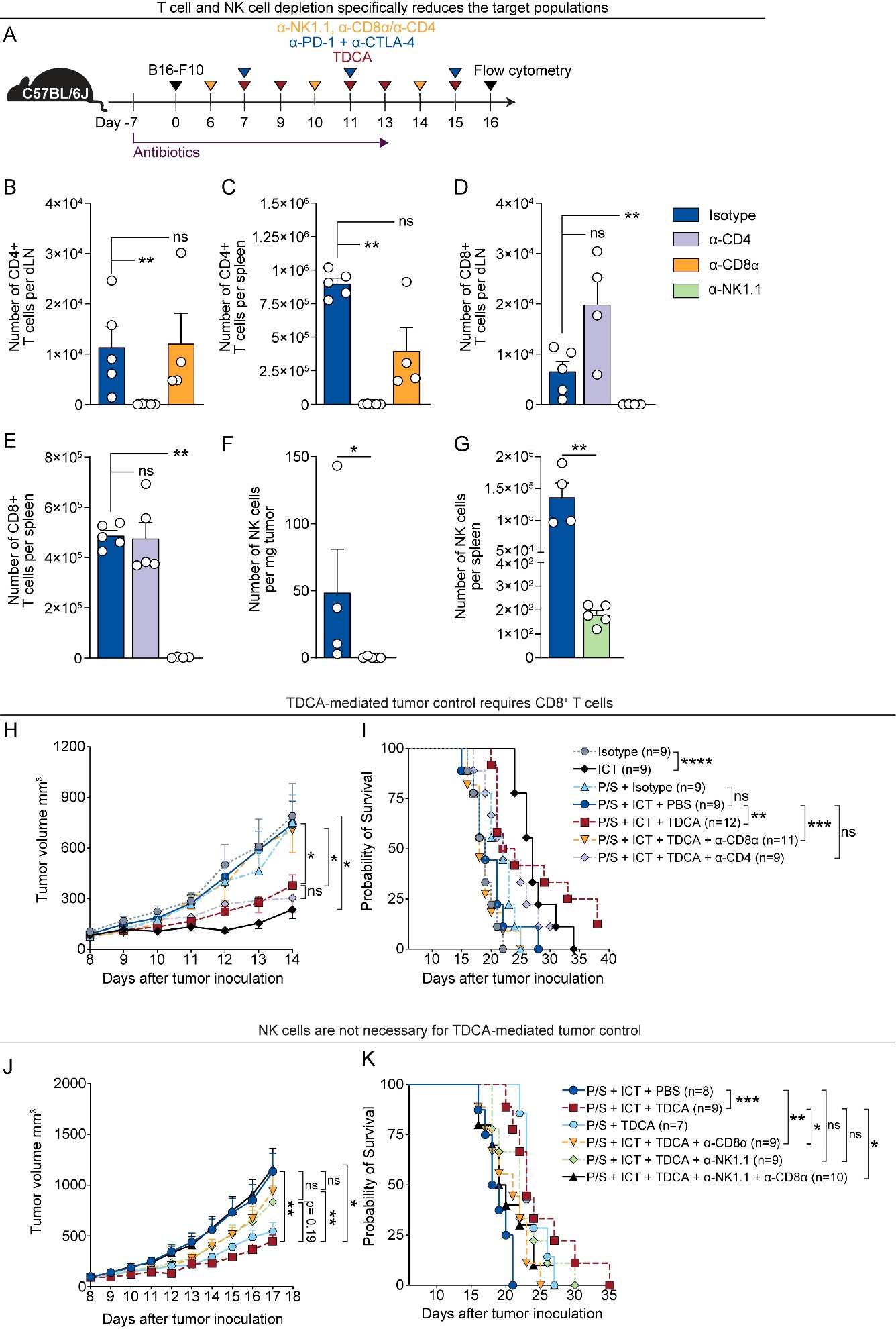

**Figure S9.** **TDCA rescue of antibiotic-induced ICT hyporesponsiveness is dependent on CD8⁺ T cells.**

**(A)** Schematic overview of CD4^+^/CD8^+^ T cell depletion in B16-F10 tumor model. Mice received penicillin/ streptomycin (0.95 g/L; 2 g/L) in drinking water throughout the experiment. 1 x 10^5^ B16-F10 cells were injected subcutaneously into the right inguinal flank. α-CD4 (100 μg), α-CD4 (100 μg) or α-NK1.1 (250 μg) intraperitoneal injections were started one day before ICT treatments. ICT or isotype control treatments began when tumor volumes reached 80 ± 20 mm³. TDCA (50 μl of 4 mM) was intratumorally injected according to the schema.

**(B-C)** Number of CD4⁺ T cells in dLNs or spleens with CD4⁺ T cell or CD8⁺ T cell depletion.

**(D-E)** Number of CD8⁺ T cells in dLNs or spleens with CD4⁺ T cell or CD8⁺ T cell depletion.

**(F-G)** Number of NK cells in tumors or spleens with NK cell depletion.

**(H)** Tumor volumes of control, CD8^+^ T cell– or CD4^+^ T cell–depleted mice from (A); n =9–12 mice per group

**(I)** Survival of tumor-bearing control, CD8^+^ T cell– or CD4^+^ T cell–depleted mice from (A); n =9–12 mice per group.

**(J)** Tumor volumes of control, CD8^+^ T cell– or NK cell–depleted mice from (A); n =7–10 mice per group.

**(K)** Survival of tumor-bearing control, CD8^+^ T cell– or NK cell–depleted mice from (A); n =7–10mice per group.

dLN, draining lymph node; NK cell, natural killer cell. Statistical analysis by one-way ANOVA, Kruskal–Wallis test (B-E, H, J), Mann–Whitney test (F-G) or Log-rank test for (I, K). Points represent individual mice. Bars denote mean ± SEM. All results are representative of ≥2 independent experiments. *p< 0.05; ****p** < 0.005; *****p** < 0.001; ns, not significant.

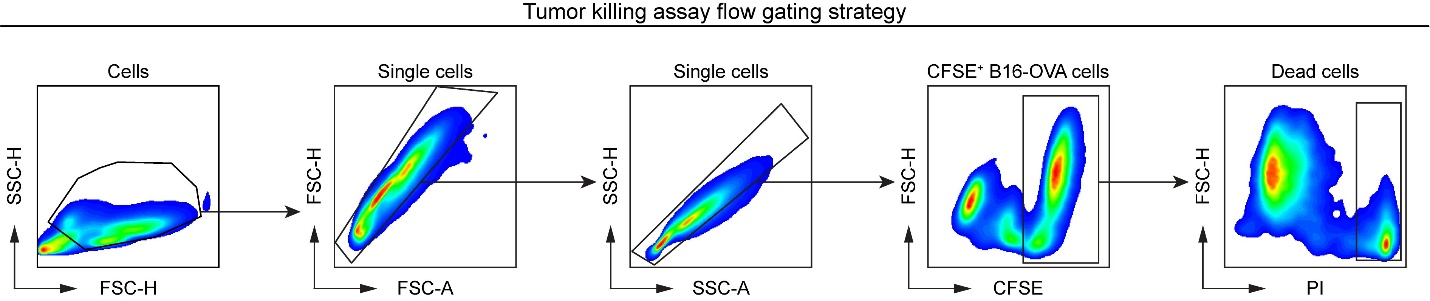

**Figure S10. Flow cytometry gating strategy for *in vitro* tumor killing assay.**
